# Animal evolution coincides with a novel degree of freedom in exocytic transport processes

**DOI:** 10.1101/591974

**Authors:** Martin Kollmar, Tobias Welz, Felix Straub, Noura Alzahofi, Klas Hatje, Deborah A. Briggs, Annette Samol-Wolf, Pawel Burkhardt, Alistair Hume, Eugen Kerkhoff

## Abstract

Exocytic transport of transmembrane receptors and secreted ligands provides the basis for cellular communication in animals. The RAB8/RAB3/RAB27 trafficking regulators function in transport processes towards the cell membrane. The small G-proteins recruit a diversity of effectors that mediate transport along microtubule and actin tracks, as well as membrane tethering and fusion. SPIRE actin nucleators organise local actin networks at exocytic vesicle membranes. By complex formation with class-5 myosins, vesicle transport track generation and motor protein activation are coordinated. Our phylogenetic analysis traced the onset of SPIRE function back to the origin of the Holozoa. We have identified SPIRE in the closest unicellular relatives of animals, the choanoflagellates, and the more distantly related ichthyosporeans. The discovery of a SPIRE-like protein encoding a KIND and tandem-WH2 domains in the amoebozoan *Physarum polycephalum* suggests that the SPIRE-type actin nucleation mechanism originated even earlier. Choanoflagellate SPIRE interacts with RAB8, the sole choanoflagellate representative of the metazoan RAB8/RAB3/RAB27 family. Major interactions including MYO5, FMN-subgroup formins and vesicle membranes are conserved between the choanoflagellate and mammalian SPIRE proteins and the choanoflagellate *Monosiga brevicollis* SPIRE protein can rescue mouse SPIRE1/2 function in melanosome transport. Genome duplications generated two mammalian *SPIRE* genes (*SPIRE1* and *SPIRE2*) and allowed for the separation of SPIRE protein function in terms of tissue expression and RAB GTPase binding. SPIRE1 is highest expressed in the nervous system and interacts with RAB27 and RAB8. SPIRE2 shows high expression in the digestive tract and specifically interacts with RAB8. We propose that at the dawn of the animal kingdom a new transport mechanism came into existence, which bridges microtubule tracks, detached vesicles and the cellular actin cytoskeleton by organising actin/myosin forces directly at exocytic vesicle membranes. The new degree of freedom in transport may reflect the increased demands of the sophisticated cellular communications in animals.

## Introduction

Animals consist of a network of specialized, interconnected tissues to increase the feeding efficiency over a single cell organism (Brunet and King 2017; Cavalier-Smith 2017). The beneficial food supply is achieved by a signalling network, enabling animals to coordinate the functional allocation of their distinct tissues. In this respect, the high energy consuming vertebrate brains, which direct senses, behaviour and reproduction, represent a sophisticated steering device (Moroz and Kohn 2016). In multicellular animals secreted factors, which interact with receptors of recipient cells, direct the interconnecting signalling machinery between cellular networks and tissues. The mammalian hypothalamus, pituitary and adrenal gland (HPA) axis controls the mammalian stress response and serves as an example of a systemic signalling system (Arnett, et al. 2016). Intracellular transport processes mediated by components of the cellular cytoskeleton facilitate the release of the signalling factors and the membrane integration of receptors. The signalling factors and the receptors travel on routes of the exocytic pathway. They are sorted at the trans-Golgi network into vesicles and are subsequently transported by motor proteins along microtubule and actin filament tracks to the cell periphery (Stenmark 2009).

In animal cells the highways and local roads model describes the cooperative transport of cargo along microtubule and actin tracks (Hammer and Sellers 2011; Hume and Seabra 2011). Microtubules enable long and fast transport (highways) and the dynamic actin cytoskeleton provides slower but very flexible end delivery of the cargo (local roads). In plants and fungi actin filaments mediate both, long and fast transport processes, as well as the final delivery (Bretscher 2003; Geitmann and Nebenfuhr 2015). In contrast to animal cells where we see a cooperative microtubule/actin transport, in plants transport functions of the actin and microtubule cytoskeletons are separated (Hammer and Sellers 2011; Geitmann and Nebenfuhr 2015). Class-11 and class-5 myosin motor proteins are the major actin based transporters in plants (MYO11), fungi (yeast MYO2p, MYO4p) and animal cells (MYO5A, 5B, 5C), respectively (Hammer and Sellers 2011; Tominaga and Nakano 2012). According to their different modes of action, the animal class-5 myosin motors slide along the actin filaments by nearly an order of a magnitude slower than their plant and yeast homologs (filament sliding velocity: vertebrate MYO5 < 1 μm/s, plant/yeast MYO11/Myo2p > 4 μm/s)(Reck-Peterson, et al. 2001; Tominaga, et al. 2003; Krementsov, et al. 2004).

RAB family GTPases are key regulators of intracellular transport processes (Stenmark 2009). Within the over 60 human RAB proteins the GTPases of the RAB8/RAB27/RAB3 branch function in the regulation of exocytic transport processes towards the plasma membrane (Klopper, et al. 2012; Rojas, et al. 2012; Varoqueaux and Fasshauer 2017). In a cascade of interweaving steps, the small GTPases target microtubule and actin motor proteins as well as membrane tethering/fusion factors to vesicles (Fukuda 2013). Several effectors of the RAB3/RAB27/RAB8 GTPases have a common RAB interaction domain consisting of the a FYVE-type zinc finger flanked by a-helical regions (SLP homology domain, SHD)(Ostermeier and Brunger 1999; Kuroda, et al. 2002; Kukimoto-Niino, et al. 2008; Fukuda 2013). Among these factors are the synaptic active zone scaffolding proteins RIM1 and RIM2, but also the myosin-5A (MYO5A)/F-actin interacting MYRIP (SLAC2-C) and melanophilin (MLPH, SLAC2-A) proteins. Sequence homology studies recently grouped the SPIRE actin nucleators into the SHD-family of RAB3/27/8 effectors (Alzahofi et al., under revision). Mammalian SPIRE1 interacts with RAB27A in a GTP dependent manner.

SPIRE family actin nucleation factors (mammalian SPIRE1 and SPIRE2) coordinate actin filament assembly in cooperation with FMN-subfamily formins (mammalian FMN1 and FMN2) at vesicle membranes (Pfender, et al. 2011; Schuh 2011)(Alzahofi et al., under revision). By complex formation with class-5 myosins, SPIRE coordinates actin filament track generation and motor protein activation (Pylypenko, et al. 2016). The RAB27A regulated recruitment of SPIRE1, FMN1, and myosin-5A (MYO5A) to vesicles has an essential function in melanosome dense core vesicle transport towards the plasma membrane (Alzahofi et al., under revision). In addition, SPIRE1, SPIRE2, FMN2 and MYO5B direct cortical transport of RAB11 vesicles in mouse oocytes (Pfender, et al. 2011; Schuh 2011). In neuronal networks a function of SPIRE, FMN and MYO5 proteins has been associated with memory and learning (Wang, et al. 2008; Wagner, et al. 2011; Pleiser, et al. 2014; Agis-Balboa, et al. 2017). Choanoflagellates are the closest unicellular relatives of animals (Carr, et al. 2008; King, et al. 2008; Ruiz-Trillo, et al. 2008). Their genomes encode proteins of the neuronal signalling machinery, such as voltage gated sodium and calcium channels (Cai 2008; Liebeskind, et al. 2011; Gur Barzilai, et al. 2012), synaptic scaffolding proteins (PSD95, Homer, Shank) and neurosecretory proteins (SNAREs and Munc18), which are part of the vesicle release machinery (Alie and Manuel 2010; Burkhardt, et al. 2011; Burkhardt and Sprecher 2017). The latter localise to the apical membrane of the choanoflagellates and may mediate the release of molecules from vesicles (Hoffmeyer and Burkhardt 2016). The discovery of the neuronal signalling proteins in choanoflagellates raises the interesting question whether this novel protein machinery was the prerequisite for the evolution of sophisticated networks of animal cell communication.

By analysing the evolution of SPIRE actin nucleators we here show that next to the exocytic membrane vesicle fusion machinery also the generation of local actin/myosin networks at vesicle membranes, as part of the cytoplasmic function of the exocytic transport pathway predates animal evolution.

## Results

### Origin of SPIRE in the ancestor of Holozoa

Actin/myosin functions in vesicle transport processes are well conserved within the family of eukaryotes (Hammer and Sellers 2011; Tominaga and Nakano 2012). The class-5 myosin motor proteins, which act as processive motors, transport cargo along actin filaments in plants, fungi and animals. In mammals there are three class-5 myosin motors (MYO5A, B, C), which all interact with SPIRE actin nucleators (Pylypenko, et al. 2016). The coordinated actin nucleation/motor protein activation by SPIRE/MYO5 has up to now only been studied in vertebrates.

To determine the origin of SPIRE and its distribution in extant species we used TBLASTN to search in sequenced and assembled eukaryotic genomes and transcriptomes taking the well-known *Drosophila melanogaster* and human SPIRE homologs as starting sequences. SPIRE proteins were identified in Ichthyosporea, Choanoflagellida and Metazoa suggesting that SPIRE is at least a holozoan invention (Fig. 1). In addition, a SPIRE-like protein was detected in the amoebozoan *Physarum polycephalum*, but not in other sequenced Amoebozoa. As long as more homologs from basal branches are not available we suggest naming this *P. polycephalum* homolog SPIRE-like. Choanoflagellida, Filasterea, and Ichthyosporea are the closest unicellular relatives of animals. We were not able to detect a SPIRE homolog in the filasterean *Capsaspora owczarzaki* suggesting that SPIRE was lost. SPIRE has also likely been lost in Ctenophora (comb jellies), Mesozoa (worm-like parasites of marine invertebrates), Placozoa (*Trichoplax adhaerens*, small flattened animal, simplest structure of all multicellular animals), Platyhelminthes (flat worms) and Nematoda (round worms). Outside vertebrates, *SPIRE* gene duplicates have only been identified in *Amoebidium parasiticum*, the salmon louse *Lepeophtheirus salmonis* and the sea squirt *Ciona intestinalis* (but not *Ciona savignyi*) so far.

**Figure 1.**
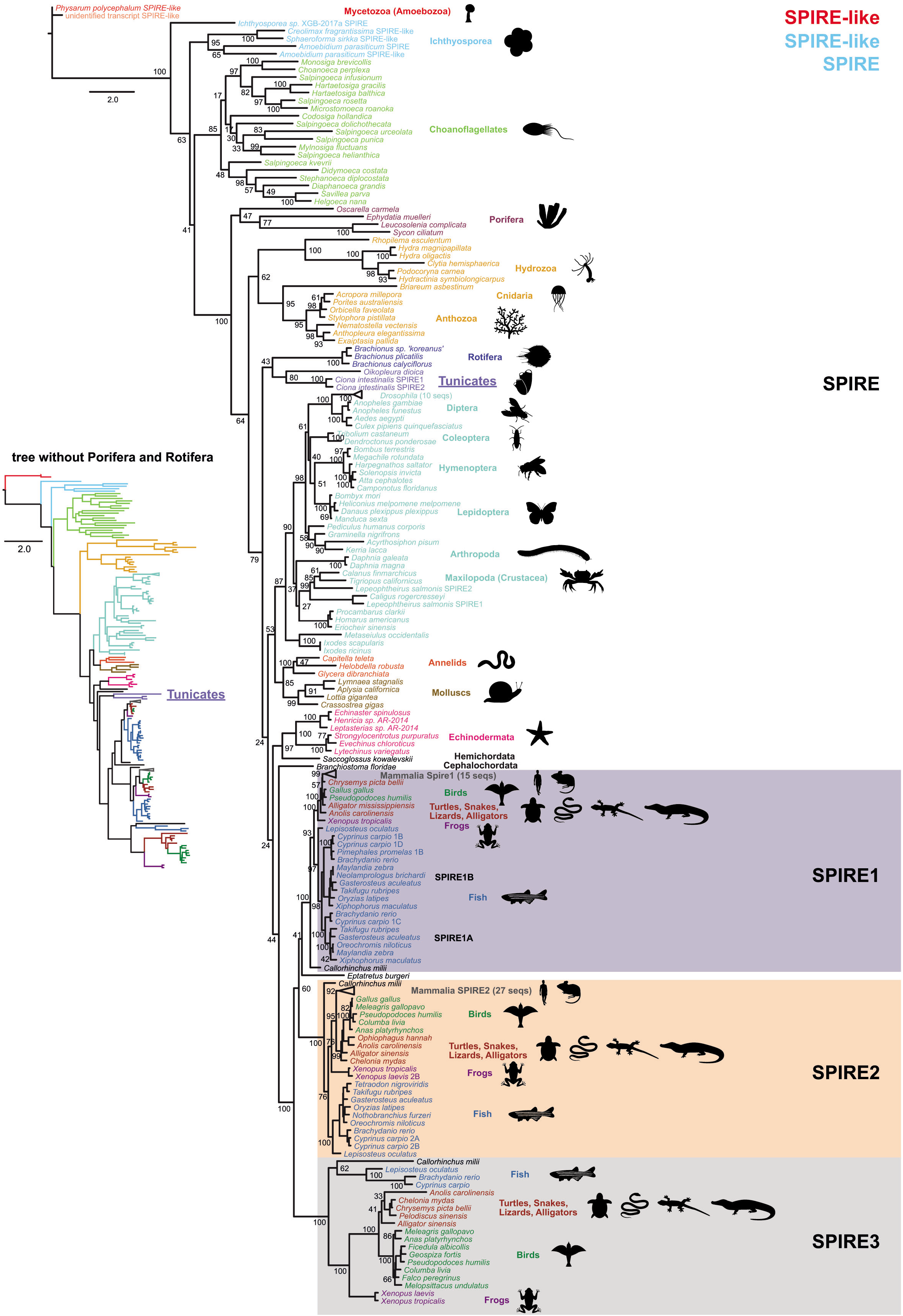
SPIRE phylogeny. IQ-TREE generated phylogeny of a CD-Hit-reduced set (90% identity level) of 217 SPIRE proteins. Support values for major branches are given as percent bootstrap replicates. For displaying purposes, mammalian and *Drosophila* SPIRE proteins have been collapsed into groups with the numbers in brackets denoting the number of sequences. Some bootstrap values at short branches have been removed for better presentation. The tree in the inlet, generated with FastTree, is based on the same data except for the four Porifera and three Rotifera SPIRE, which were removed. In this tree, the Tunicata group sister to the Vertebrata indicating some long-range attraction by the Porifera and Rotifera SPIRE sequences. The colouring of the branches is identical to the colour of species names in the large tree.

### Concordance of SPIRE and species phylogeny

The phylogenetic tree of the SPIRE proteins is in accordance with the commonly agreed tree of the Holozoa and especially the Metazoa (Pennisi 2003) with the Lophotrochozoa (Annelida and Mollusca) sister to the Arthropoda (joint-legged animals) and together forming the Protostomia, and with the Echinodermata (hedgehog skin), the Cephalochordata (lancelets) and the Hemichordata branching basal to the Vertebrata and together with these forming the Deuterostomia (Fig. 1). The respective branchings are highly supported by bootstrap values (ML analyses). The tunicate SPIRE homologs are influenced by the Porifera (sponges) and Rotifera (wheel animals). When the latter are included in the alignment, the tunicate SPIRE group outside the Bilateria, while when the sponge and rotifer SPIRE are excluded, the tunicate SPIRE group sister to the vertebrates (Fig. 1). Within the Vertebrata we identified a single *SPIRE* gene in *Eptatretus burgeri* (inshore hagfish, belongs to the jawless fish of the Cyclostomata) and three groups of SPIRE homologs in Gnathostomata (jawed vertebrates) which we termed SPIRE1 to SPIRE3 and which we labelled with respect to the long-known two human SPIRE homologs (Fig. 1). The exact grouping of the three SPIRE subtypes with respect to each other could not be resolved (only very weak bootstrap support of the respective branchings). Depending on whether the *E. burgeri* SPIRE and/or the *Callorhinchus milii* SPIRE3 (Australian ghostshark, belongs to the cartilaginous fish of Chondrichthyes) are included in the dataset, the SPIRE2 group with the SPIRE1 or the SPIRE3, respectively. While the phylogeny of the invertebrates always remains stable, the *E. burgeri* SPIRE was found both within and outside the other vertebrate SPIRE (Fig. 1). These data indicate that more cyclostomatan SPIRE sequences are needed to get a stable phylogeny of the vertebrate SPIRE duplicates. The current data can also not yet be related to the two vertebrate whole-genome duplication (WGD) events, which most likely happened before the split of the Cyclostomata and the Gnathostomata (Smith, et al. 2013). The single *E. burgeri* SPIRE suggests two SPIRE gene duplication events in the ancestor of the Gnathostomata independent of the two WGD, but as long as the SPIRE phylogeny is not entirely resolved it is also possible that *E. burgeri* lost one or more SPIRE homologs. The SPIRE3 subtype has independently been lost by the last common ancestor of the Mammalia and the last common Euteleostei (bony fish) ancestor.

### Domain array of SPIRE proteins throughout evolution

Sequence comparison revealed that all identified metazoan SPIRE homologs (except hydrozoan SPIRE and some SPIRE3) consist of an N-terminal KIND domain (kinase non-catalytic C-lobe domain)(Ciccarelli, et al. 2003; Pechlivanis, et al. 2009; Vizcarra, et al. 2011; Zeth, et al. 2011), which mediates the interaction with FMN-subgroup formins (Quinlan, et al. 2007; Pechlivanis, et al. 2009), followed by four G-actin binding WH2 motifs (WASP-homology 2) (Quinlan, et al. 2005; Dominguez 2016), a myosin-5 interaction motif (GTBM, globular tail domain binding motif) in the middle (Pylypenko, et al. 2016), and a tandem domain consisting of a so-called SB (SPIRE-box) motif and a FYVE-type zinc finger domain (named after the cysteine-rich proteins Fab 1, YOTB, Vac 1, and EEA1) at the C-terminus (Kerkhoff, et al. 2001), which contributes to vesicle membrane localisation by direct interaction with the phospholipid bilayer (Tittel, et al. 2015)(Fig. 2). The C-terminal SPIRE sequences share homology with the Slp homology domain (SHD) of exocytic transport regulators such as SLP3-A, RIM1, Rabphilin, SLAC2-A (melanophilin, MLPH), and SLAC2-C (MYRIP) (Kerkhoff, et al. 2001; Fukuda 2013). The SPIRE SHD-like sequences were recently shown to mediate the interaction of SPIRE1 with RAB27A and are essential for a functional cooperation of SPIRE/MYO5A in melanosome transport (Alzahofi et al., under revision).

**Figure 2.**
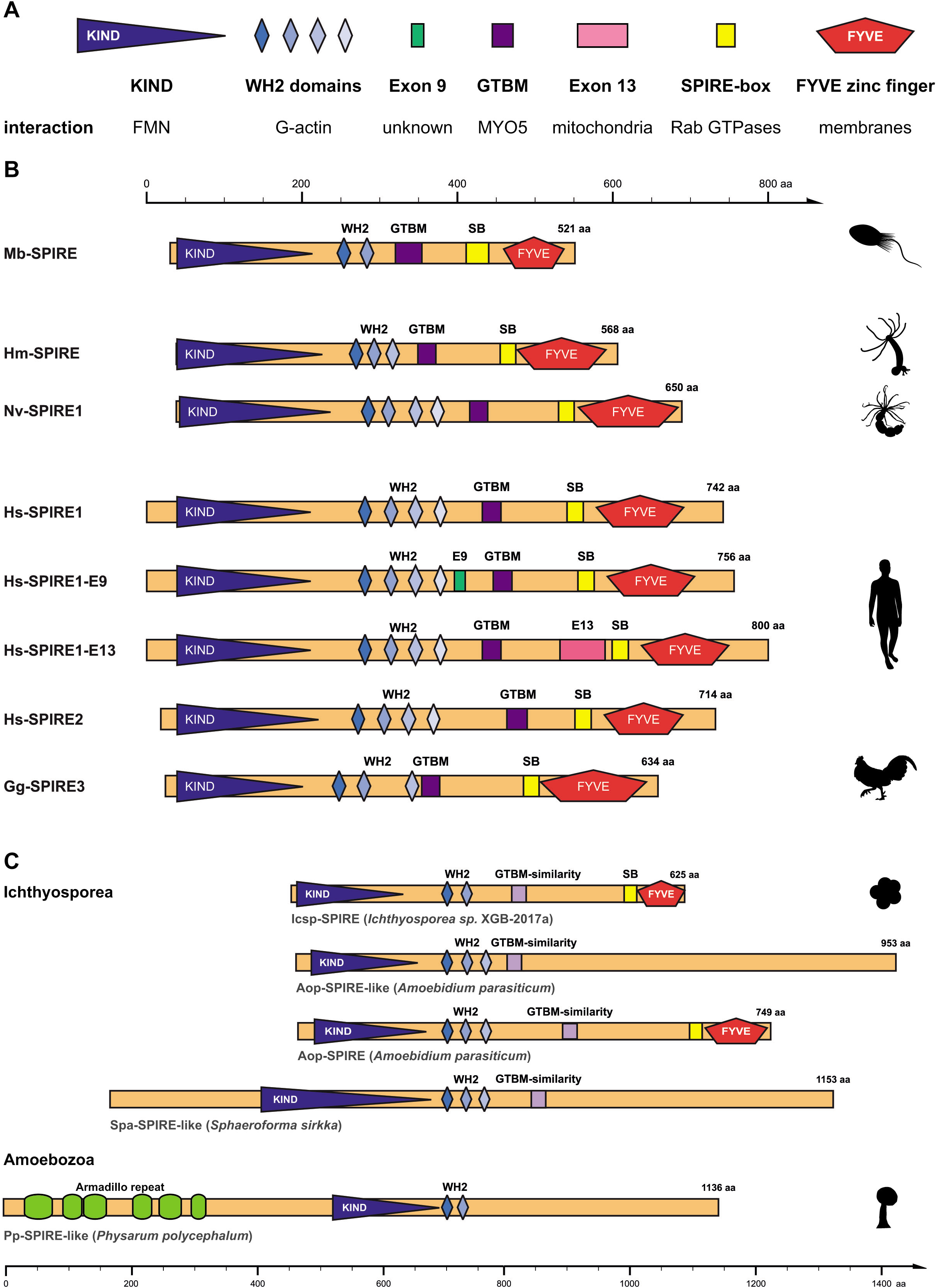
Functional SPIRE protein domains are conserved across species. (A) Overview on functional SPIRE protein domains and their specific interaction partners and intracellular structures. KIND, *kinase non-catalytic C-lobe domain;* WH2, *Wiskott-Aldrich-Syndrome protein homology 2*; GTBM, *globular tail domain binding motif;* FYVE, after Fab 1/YOTB/Vac 1/EEA1. (B) Overview of invertebrate and vertebrate SPIRE proteins and their respective domain organizations. Species abbreviations are: Mb, *Monosiga brevicollis;* Hm, *Hydra magnipapillata;* Nv, *Nematostella vectensis;* Hs, *Homo sapiens;* Gg, *Gallus gallus*. (C) The domain organization of ichtyosporean and amoebozoan SPIRE/SPIRE-like proteins are shown. Numbers indicate amino acids.

In contrast, the *P. polycephalum* SPIRE-like protein has a KIND domain and two WH2 motifs with strong homology to the other SPIRE proteins, but contains an N-terminal extension of about 500 amino acids encoding armadillo repeats (Fig. 2C). In addition, it misses the GTBM and SB motifs and the C-terminal FYVE domain, which are characteristic for all SPIRE proteins. The domain architectures of the ichthyosporean and choanoflagellate SPIRE homologs are identical to the metazoan ones except that these SPIRE encode only two or three WH2 domains and considerably differ in their proposed MYO5 golobular tail domain binding motifs (GTBMs)(Fig. 2). The sequence homology of the WH2 domains suggests that the ancient holozoan SPIRE contained three or even four WH2 domains, and that extant choanoflagellates picked individual combinations of these. The sauropsid SPIRE3 homologs do not contain the third WH2 domain or the sequence has been changed beyond recognition (Fig. 2).

### The choanoflagellate SPIRE protein interacts with myosin-5

Our genome studies identified the origin of SPIRE proteins in the last common ancestor of the Holozoa (Fig. 1). Choanoflagellates encode a single class-5 myosin motor protein (Fig. 3A), and a conserved region between the last WH2 domain and the SB motif distantly resembling the MYO5 globular tail domain binding motif (GTBM) in human SPIRE proteins could be detected in the choanoflagellate SPIRE sequences (Fig. 4C). To answer the question of the origin of SPIRE functions, we have experimentally addressed the interactions of the SPIRE protein from *M. brevicollis* (Mb-SPIRE).

**Figure 3.**
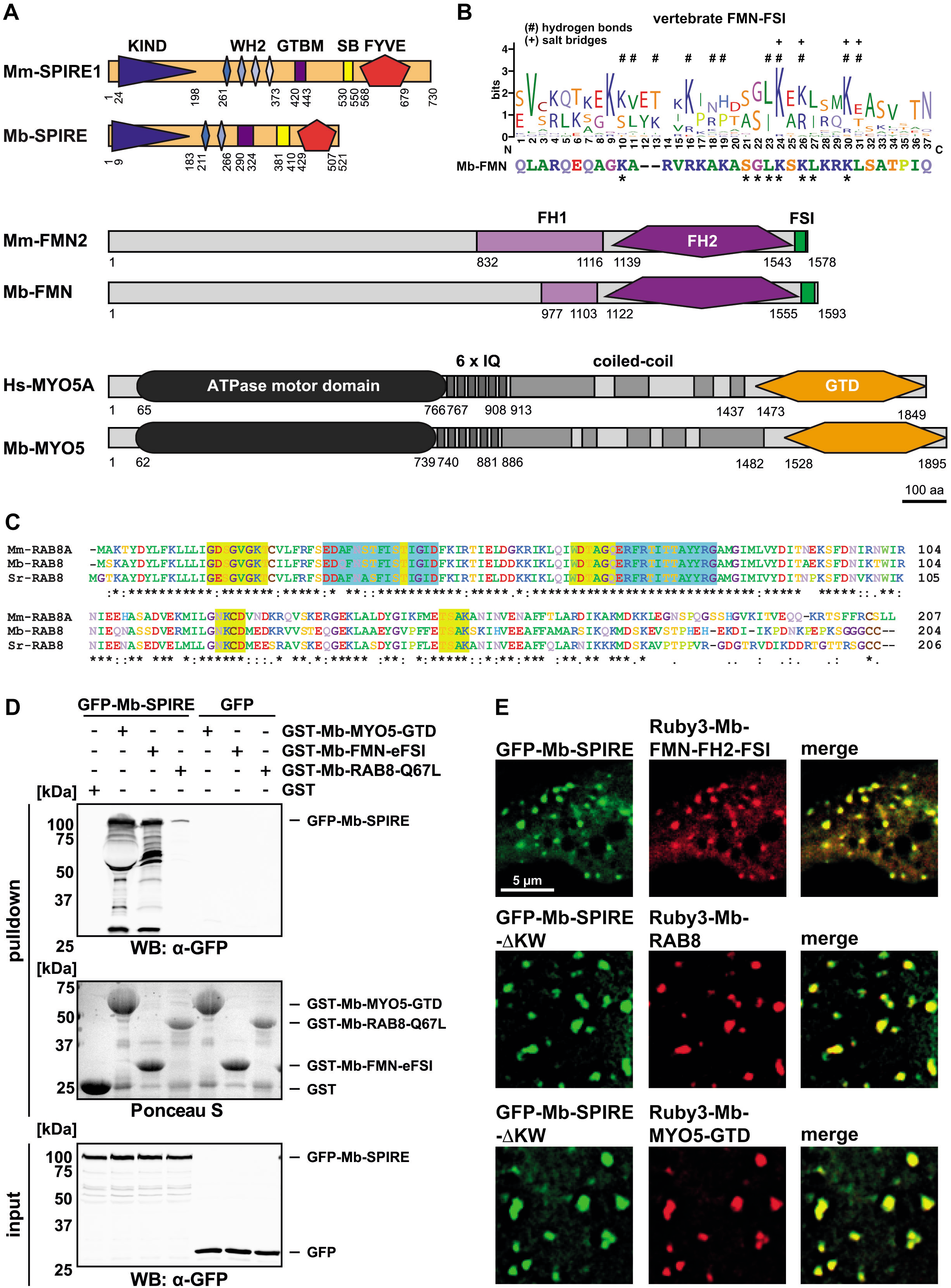
Conserved SPIRE protein interactions. (A) Domain organization of vertebrate (Hs, *Homo sapiens;* Mm, *Mus musculus*) and *Monosiga brevicollis* (Mb) SPIRE, formin subfamily formins (FMN) and myosin-5 (MYO5) proteins. SPIRE proteins share KIND (kinase non-catalytic C-lobe domain), WH2 (Wiskott-Aldrich-Syndrom protein homology 2) domains, GTBM (globular tail domain binding motif), SB (SPIRE-box) and FYVE-type zinc finger (after Fab1/YOTB/Vac1/EEA1). FMN proteins share the formin homology domains (FH1 and FH2), the C-terminal formin-SPIRE interaction sequence (FSI) and a so far uncharacterized N-terminus. MYO5 proteins share all functional domains, including the ATPase motor domain, calmodulin-binding IQ motifs, coiled-coil regions and a C-terminal cargo-binding globular tail domain (GTD). Protein structures are drawn in scale and numbers indicate amino acids. (B) WebLogo showing amino acid conservation within vertebrate FMN-FSI protein sequences. Residues forming hydrogen bonds (#) and salt bridges (+) with the SPIRE-KIND are labeled respectively. Below the WebLogo the respective C-terminal sequence part of Mb-FMN is depicted, adjusted to the WebLogo and conserved residues are labeled (*). (C) A multiple protein sequence alignment is shown for RAB8 proteins from *Mus musculus* (Mm), *Monosiga brevicollis* (Mb) and *Salpingoeca rosetta* (Sr). The highly conserved G motifs within the G domain are labeled in yellow, residues forming the interacting switch I and switch II regions are labeled in blue. (D) GST-pulldown assay with purified GST-Mb-MYO5-GTD, GST-Mb-FMN-eFSI and GTP-locked GST-Mb-RAB8-Q67L and HEK293 cell lysates transiently over-expressing full-length AcGFP1-tagged Mb-SPIRE (GFP-Mb-SPIRE). GST and GFP, respectively, were used as controls and Ponceau S staining shows equal amounts of GST-tagged proteins. N = 2 experimental repeats. (E) The localization of transiently co-expressed tagged full-length Mb-SPIRE and Mb-SPIRE-ΔKW (AcGFP1; GFP-Mb-SPIRE, GFP-Mb-SPIRE-ΔKW; green), Mb-FMN-FH2-FSI (mRuby3; Ruby3-Mb-FMN-FH2-FSI; red), Mb-RAB8 (mRuby3; Ruby3-Mb-RAB8; red) and Mb-MYO5-GTD (mRuby3; Ruby3-Mb-MYO5-GTD; red) was analyzed by fluorescence microscopy. Deconvoluted pictures indicate the localization of the proteins on vesicular structures and colocalization of GFP-Mb-SPIRE and Ruby3-Mb-FMN-FH2-FSI, GFP-Mb-SPIRE-ΔKW and Ruby3-Mb-RAB8, and GFP-Mb-SPIRE-ΔKW and Ruby3-Mb-MYO5-GTD (merge). *Scale bars* represent 5 μm. 5 cells were recorded for each condition and the cytoplasmic region of one representative cell is presented here.

**Figure 4.**
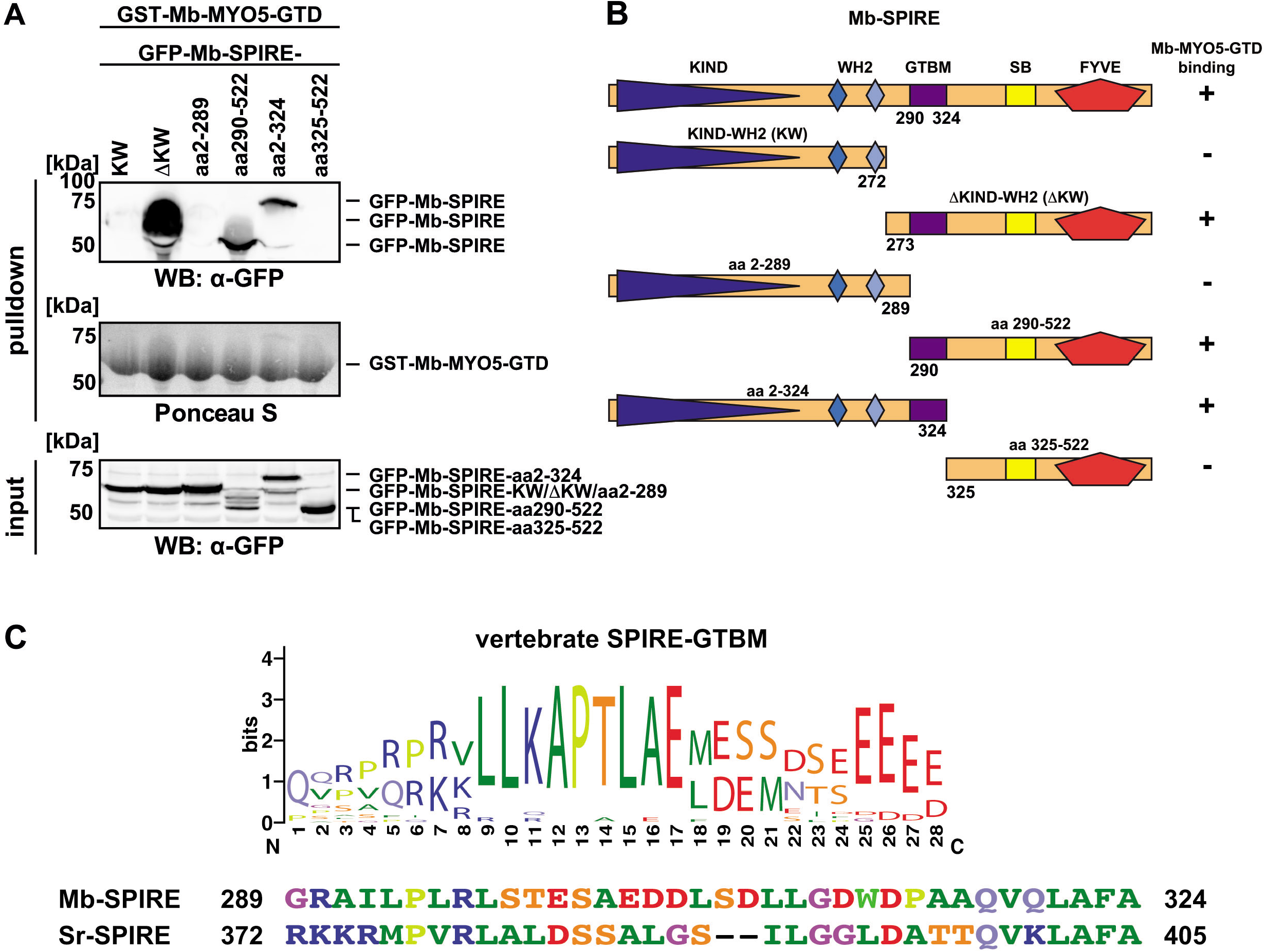
Characterization of the Mb-SPIRE Mb-MYO5 interaction sequence. (A) GST-pulldown assay with purified GST-Mb-MYO5-GTD and HEK293 cell lysates transiently over-expressing AcGFP1-tagged Mb-SPIRE (GFP-Mb-SPIRE) N-terminal and C-terminal protein fragments. Ponceau S staining shows equal amounts of GST-Mb-MYO5-GTD proteins. N = 2 experimental repeats. KW, *KIND-WH2;* WB, *Western blotting;* numbers indicate amino acids. (B) Schematic representation of N-terminal and C-terminal Mb-SPIRE protein fragments as used in (A) and their capacity to bind (+) or not bind (-) to Mb-MYO5-GTD. (C) WebLogo showing amino acid conservation within vertebrate SPIRE GTBM protein sequences. The corresponding SPIRE protein sequences of *Monosiga brevicollis* (Mb) and *Salpingoeca rosetta* (Sr) and their respective amino acid boundaries are shown below the WebLogo.

A full-length coding region of the *Mb-SPIRE* cDNA was generated by gene synthesis. We also synthesised a cDNA encoding the globular tail domain of the *M. brevicollis* myosin-5 motor protein (Mb-MYO5)(Fig. 3A). In GST-pulldown assays a purified bacterially expressed GST-Mb-MYO5-GTD fusion protein strongly interacts with AcGFP-tagged full-length Mb-SPIRE (GFP-Mb-SPIRE), which was transiently expressed in HEK293 cells (Fig. 3D). The AcGFP-Mb-SPIRE protein does not interact with GST nor does the AcGFP protein interact with GST-Mb-MYO5-GTD. Further support for an interaction was obtained from colocalisation studies of transiently expressed choanoflagellate SPIRE and MYO5 proteins in human HeLa cells. The choanoflagellate N-terminal deletion mutant AcGFP-Mb-SPIRE-ΔKW colocalises with mRuby3 tagged Mb-MYO5-GTD at vesicular structures (Fig. 3E). By deletion mutation of the AcGFP-Mb-SPIRE protein we mapped the Mb-MYO5 interaction sequences to a sequence of 36 amino acids the central region of the Mb-SPIRE protein, in between the WH2 domain and the SPIRE-box, which is exactly the region where the MYO5 globular tail domain binding motif (GTBM) of the mammalian SPIRE proteins is located (Fig. 4A, B). Comparison of the Mb-SPIRE MYO5 interaction motif with vertebrate SPIRE GTBM motifs shows only very low sequence homology (Fig. 4C). However, the array of basic (blue) /hydrophobic (green) /acidic residues (red) in the vertebrate GTBM might be conserved in the Mb-SPIRE GTBM (Fig. 4C). Comparing their sequences, the Mb-MYO5 globular tail domain is closely related to the vertebrate MYO5-GTDs (Pylypenko, et al. 2013; Kollmar and Muhlhausen 2017) and it remains to be analysed whether Mb-SPIRE contacts Mb-MYO5 in a similar way as it was found in mammals. The strong interaction of Mb-SPIRE and Mb-MYO5, however, indicates that the cooperation of actin nucleation and motor protein activation is conserved between choanoflagellates and mammals. Ichthyosporean SPIRE proteins encode sequences very similar to the choanoflagellate SPIRE-GTBMs (Fig. 2C, bright purple). However a potential ichthyosporean SPIRE/MYO5 interaction remains to be analysed.

### Formin/SPIRE interaction in choanoflagellates

Another hallmark of SPIRE proteins is their interaction with FMN-subgroup formins (*Drosophila* Cappuccino, mammalian FMN1 and FMN2) (Quinlan, et al. 2007; Pechlivanis, et al. 2009). In mouse and also in fly oocytes SPIRE and FMN-subgroup formins cooperate in generating an ooplasmic actin meshwork (Dahlgaard, et al. 2007; Pfender, et al. 2011). At mouse melanosome membranes, SPIRE1 and FMN1 organise a local actin network necessary to spread the melanosomes by MYO5A motor protein function (Alzahofi et al., under revision). The cooperation of the two actin nucleators is mediated by a direct interaction of the SPIRE-KIND domain and the C-terminal formin SPIRE interaction (FSI) motif (Pechlivanis, et al. 2009; Vizcarra, et al. 2011; Zeth, et al. 2011). Considering the high similarity of the formin homology domains (FH1, FH2), the short FSI motif, which is characterised by a cluster of basic residues (Pechlivanis, et al. 2009; Schonichen and Geyer 2010)(Fig. 3B), represents the only obvious defining feature of FMN subgroup formins. We could identify formins in Choanoflagellida and Ichthyosporea that encode FSI-like motifs at their C-terminal ends (Fig. 3A, B, data not shown). We have fused extended sequences of the putative choanoflagellate FSI motif to GST (GST-Mb-FMN-eFSI) and have performed GST-pulldown experiments with AcGFP-Mb-SPIRE transiently expressed in HEK293 cells. The GST-Mb-FMN-eFSI protein strongly interacts with AcGFP-Mb-SPIRE, but not with AcGFP alone (Fig. 3D). In addition, transient expression studies in human HeLa cells shows a colocalisation of an mRuby3-tagged C-terminal Mb-FMN protein (mRuby3-FMN-FH2-FSI) and AcGFP-Mb-SPIRE at vesicular structures (Fig. 3E).

In summary our GST-pulldown and colocalisation studies show that major features such as myosin-5 and FMN-subgroup formin interactions are conserved between mammals and choanoflagellates. In addition, the vesicular localisation of the choanoflagellate Mb-SPIRE in human HeLa cells further shows that the choanoflagellate SPIRE protein is targeted to vesicular membranes, just as it was found for its mammalian homologs.

### Choanoflagellate SPIRE is a RAB8 effector

The melanosome localisation of mouse SPIRE1 is facilitated by the interaction of C-terminal SPIRE1 sequences (SPIRE-box, FYVE-type, C-terminal flanking sequences) with RAB27 (Alzahofi et al., under revision). SPIRE1 shares its RAB27 interaction domain with a group of proteins, which encode the central FYVE-type zinc finger motif and flanking helical regions (SHD RAB27 effectors)(Kuroda, et al. 2002; Fukuda 2013). The SHD RAB27 effectors are involved in different steps of the secretory transport pathway. The transition to multicellularity in animals saw an increased need of cargo sorting (Varoqueaux and Fasshauer 2017). This is reflected in an increase of trafficking routes, which requires an in parallel amplification of the regulatory molecular machinery. Genome studies show that the mammalian RAB GTPase family regulators of the exocytic/secretory pathway, RAB27, RAB3 and RAB8, all originate from a RAB8 GTPase gene in the last eukaryotic common ancestor (LECA)(Klopper, et al. 2012; Varoqueaux and Fasshauer 2017). In the *M. brevicollis* genome we identified a single *RAB8* gene, Mb-RAB8, whose encoded protein sequence has high homology to mammalian RAB8 in the G domain, but is quite different in the C-terminal sequences (Fig. 3C). The G domain of small GTPases interacts with effector proteins, whereas the C-terminal sequences direct the GTPase to specific vesicle populations (Pylypenko, et al. 2018). Cysteine residues at the C-terminal ends of RAB GTPases are prenylated and the hydrophobic post-translational modifications anchor the RAB proteins in phospholipid bilayers (Pylypenko, et al. 2018). The choanoflagellate RAB8 proteins from *M. brevicollis* and *S. rosetta* encode two C-terminal cysteines, whereas the mammalian RAB8 proteins have only one (Fig. 3C).

In analogy to the mammalian SPIRE1/RAB27 interaction we here have analysed the interaction of Mb-SPIRE with Mb-RAB8 by GST-pulldown assays. A purified bacterially expressed GST-fusion protein of the Mb-RAB8-Q67L mutant (GST-Mb-RAB8-QL, GTP-locked form), pulls transiently expressed AcGFP-Mb-SPIRE from HEK293 lysates (Fig. 3D). GST-Mb-RAB8-QL mutant does not interact with AcGFP alone. We also analysed the localisation of an mRuby3-tagged choanoflagellate Mb-RAB8 protein (mRuby3-Mb-RAB8) in human HeLa cells and found it strongly colocalising with the mouse RAB8A protein (data not shown). When transiently co-expressed with AcGFP-tagged C-terminal *M. brevicollis* SPIRE protein (AcGFP-Mb-SPIRE-ΔKW), we gained further support for complex formation of Mb-SPIRE and Mb-RAB8 by finding them highly colocalising at vesicular structures (Fig. 3E). Considering the high conservation of vertebrate RAB8 function in vesicle transport processes towards the cell membrane, the choanoflagellate SPIRE/RAB8 interaction shown here suggests that that SPIRE proteins originated as part of the basic exocytic transport machinery.

### Choanoflagellate Mb-SPIRE rescues mouse SPIRE1/2 function in melanosome transport

Mammalian SPIRE/FMN actin nucleation mediates MYO5 actin motor protein dependent transport of RAB11 vesicles in mouse oocyte and melanosome dense core vesicle transport in mouse melanocytes (Pfender, et al. 2011; Schuh 2011)(Alzahofi et al., under revision). In both cases the actin nucleators cooperate in the generation of an actin meshwork at vesicle membranes. Neither SPIRE nor the FMN formin alone can nucleate the local actin networks. Knockout of FMN1 or knockdown of SPIRE1/2 in mouse melanocytes causes a clustering of the melanosomes around the cell nucleus (perinuclear clustering) (Alzahofi et al., under revision)(Fig. 5). The phenotype is also found when RAB27, MYO5A or melanophilin functions are inhibited (Alzahofi et al., under revision)(Hume and Seabra 2011). Just as the F-actin binding protein melanophilin, SPIRE1 is an effector of RAB27 and RAB27 targets SPIRE1 towards melanosomes (Alzahofi et al., under revision). Transgenic expression of an eGFP-tagged SPIRE1 protein can rescue the SPIRE1/SPIRE2 double knockdown and disperse the melanosomes in the cytoplasm similar to the wild-type situation (Fig. 5). Transgenic expression of eGFP alone had no effect on melanosome localisation. Also eGFP-SPIRE2 can rescue the perinuclear clustering phenotype, however here we see in several cells a hyper-dispersion, in which the melanosomes accumulate close to the cell membrane (Fig. 5A, B).

**Figure 5.**
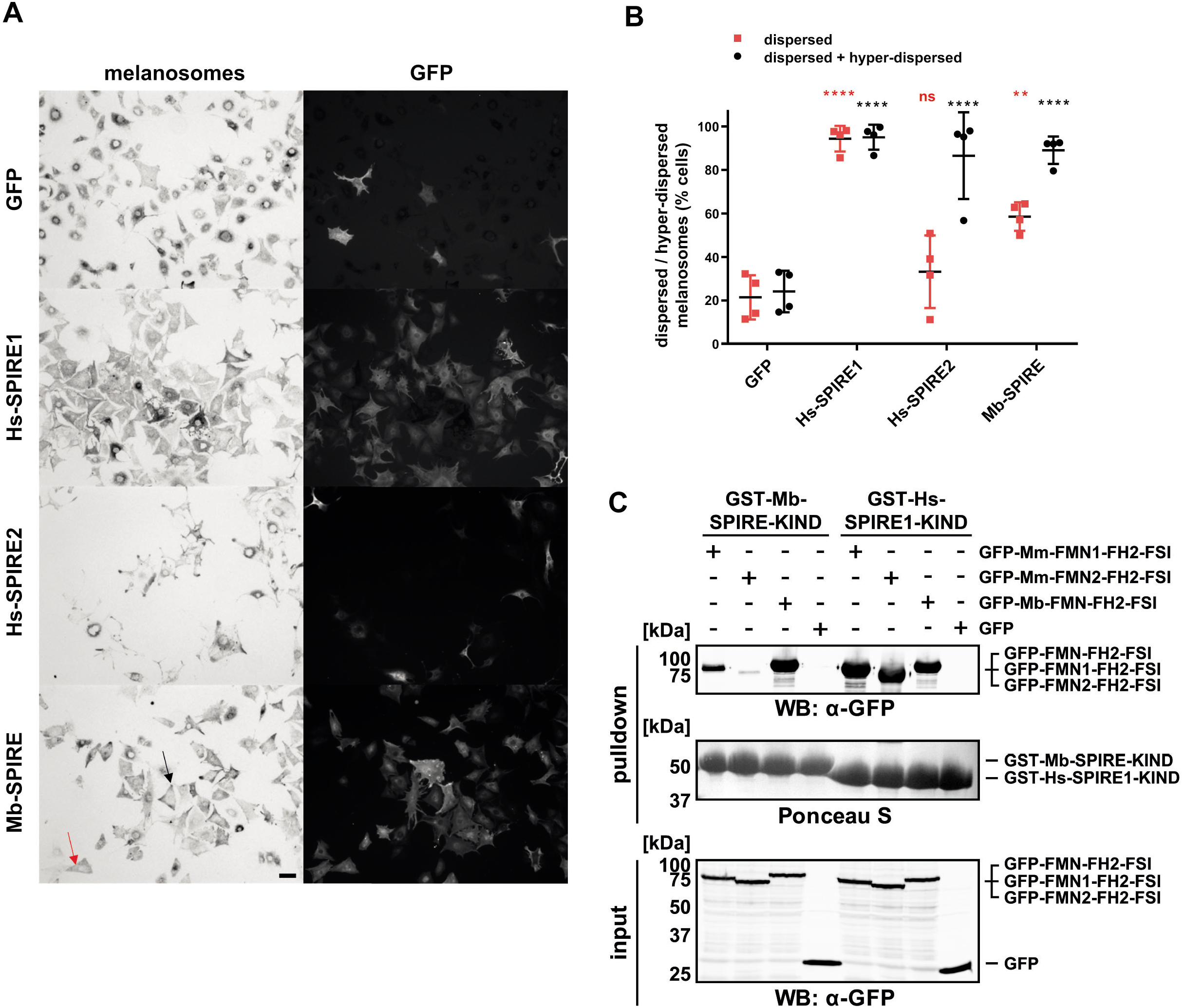
Expression of choanoflagellate (*Monosiga brevicollis*) Mb-SPIRE protein rescues melanosome dispersion in mouse melanocytes depleted of endogenous SPIRE1/2. Melan-a cells (wild-type melanocytes) were depleted of endogenous SPIRE1/2 by siRNA transfection and 72 hours later infected with adenoviruses expressing the indicated proteins. Cells were fixed 24 hours later, processed for immunofluorescence and imaged using bright-field and fluorescence optics to observe melanosome and protein distribution/expression (see experimental procedures). (A) Example images showing distribution of melanosomes and GFP in fields of cells expressing each protein. *Scale bar* represents 100 μm. Red and black arrows highlight examples of cells in which melanosomes are classed as dispersed or hyper-dispersed. (B) is a scatter plot showing the percentage of melanocytes in each population in which melanosome distribution is classed as dispersed and hyper-dispersed. Each data point indicates the percentage of cells in each class in each of 3 independent experiments and represents the average percentage of cells with the indicated phenotype in five fields of view. ****, ** and ns indicate significant differences p = < 0.0001, < 0.01 and > 0.05 compared with GFP expressing cells as determined by one-way ANOVA. Asterisk colour indicates the class of data tested. No significant differences were seen between other datasets and GFP or KW-RAB27A. Bars indicate the mean and 25^th^ and 75^th^ percentile of data. (C) The cross-species interaction of the *M. brevicollis* and mammalian SPIRE and FMN proteins was analysed by GST-pulldown experiments. The SPIRE KIND domains were fused to GST and purified from bacteria. The FMN-FH2-FSI proteins were transiently expressed as GFP fusion proteins in HEK293 cells. Western blots (WB) of the loaded and pulled proteins and a Ponceau S stain of the GST-KIND proteins are shown.

To test whether the SPIRE/FMN cooperation in actin nucleation is evolutionary conserved between choanoflagellates and mammals, we have analysed whether the *M. brevicollis* Mb-SPIRE protein can rescue the SPIRE1/SPIRE2 function in mouse melanocytes. Transgenic expression of eGFP-Mb-SPIRE rescues the perinuclear clustering of SPIRE1/SPIRE2 knockdown in mouse melanocytes. Interestingly Mb-SPIRE causes a significant hyper-dispersion phenotype, similar to the mouse SPIRE2 rescue. In agreement with the rescue of the perinuclear melanosome clustering, we found an interaction of choanoflagellate Mb-SPIRE with the mouse FMN1-FH2-FSI protein in GST-pulldown assays (Fig. 5C), suggesting that Mb-SPIRE cooperates with mouse FMN1 to generate local actin filament tracks for melanosome dispersion.

The nature of the hyper-dispersion phenotype remains speculative. SPIRE2 however preferentially interacts with RAB8 and only very weakly with RAB27 (Fig. 7). Choanoflagellate RAB8 and mouse RAB8 are very similar (Fig. 3C) and a preferential interaction of Mb-SPIRE with mouse RAB8 could be a reason for the observed hyper-dispersion. This however should be addressed in future experiments.

### Mammalian specification of SPIRE function

Genome duplications generated three *SPIRE* genes in vertebrates (*SPIRE1, SPIRE2, SPIRE3*), two of which are preserved in mammals (*SPIRE1* and *SPIRE2;* Fig. 1). The genes of the three vertebrate SPIRE subfamilies have identical exon-intron gene structures except for the exons encoding the highly variable region between the GTBM and SB motifs (Fig. 6A, data not shown). The third exon in the *SPIRE2* homolog corresponds to the fusion of exon 3 and exon 4 of SPIRE1/SPIRE3 and is most likely the result of an intron-loss event (Fig. 6A). *SPIRE1* homologs contain differentially included exons before the GTBM and the SB motifs (Fig. 6A). These exons are not present in the single hagfish *E. burgeri SPIRE* homolog, which might support its ancestry before duplication of the ancient gnathostomian *SPIRE* homolog. The function of the alternative *SPIRE1* exon 9 (E9) is currently unknown. The alternative exon 13 (E13) targets SPIRE1 function towards mitochondria (Manor, et al. 2015). Our PCR-gene expression analysis revealed that the two alternatively spliced exons are not expressed in the same mRNA (Fig. 6C). This results in four SPIRE proteins in mammals, three SPIRE1 variants and a single SPIRE2 protein (Fig. 2, Fig. 6B). Cellular localisation studies by transient expression of Myc-epitope- and AcGFP-tagged mammalian SPIRE proteins in HeLa cells shows a vesicular localisation of SPIRE1, SPIRE1-E9 and SPIRE2 and a mitochondrial localisation of SPIRE1-E13 (also denoted as SPIRE1C before, (Manor, et al. 2015))(Fig. 6E). Colocalisation studies of the vesicular SPIRE proteins revealed a nearly identical localisation of SPIRE1 and SPIRE1-E9 and a significant overlap in localisation of SPIRE1 and SPIRE2 (Fig. 6E). Specifically, vesicles close to the cell boarder are enriched in SPIRE2 expression. SPIRE1-E13 in contrast colocalises with the mitotracker mitochondria marker dye and was not detected on transport vesicles (Fig. 6E).

**Figure 6.**
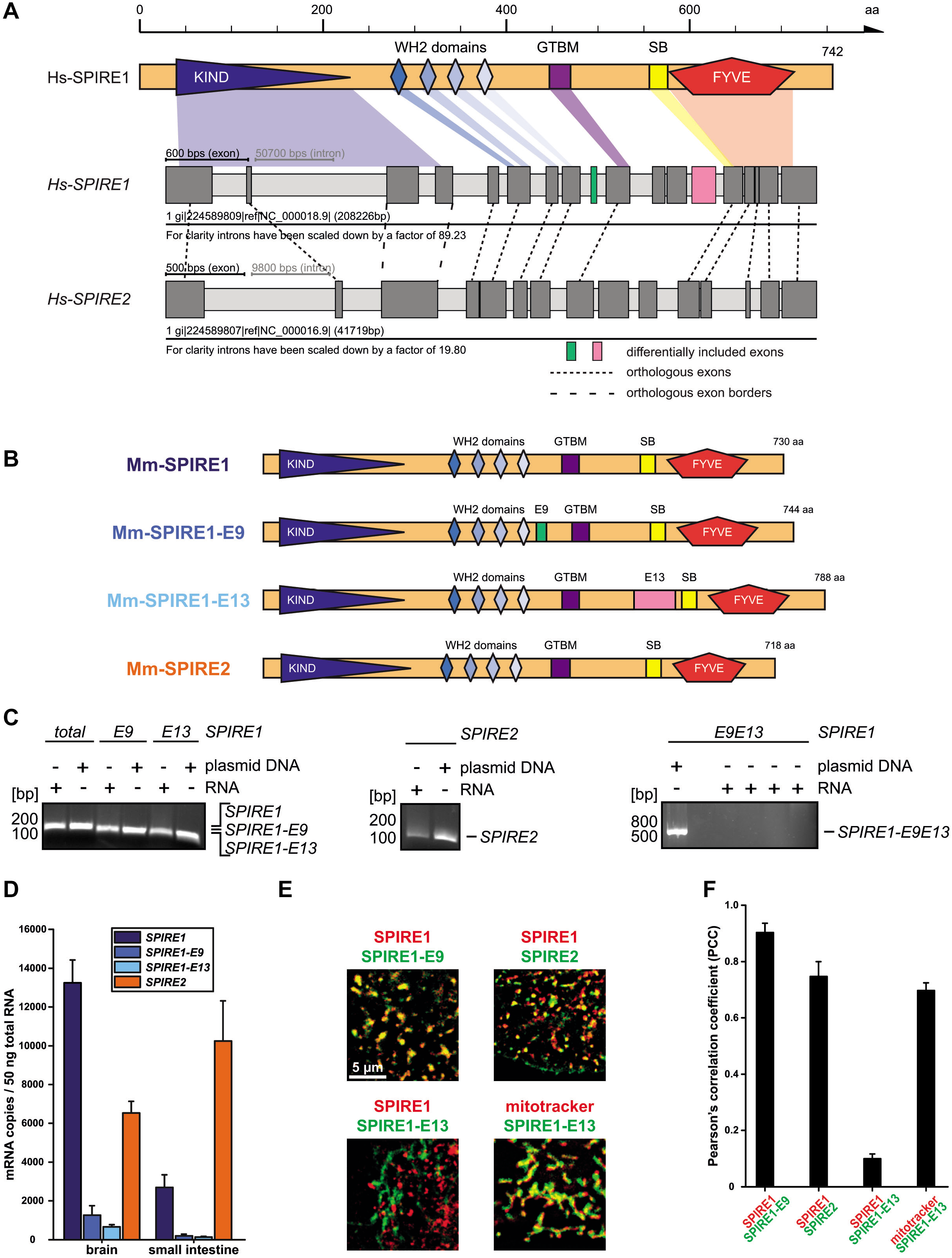
Conserved SPIRE genes show different expression patterns in distinct tissues and differences in protein localization. (A) Schematic representation of the human SPIRE1 protein and the predicted gene structure of human *SPIRE1* and *SPIRE2* genes. Conserved SPIRE protein domains and motifs are indicated and associated to their respective exon sequences. Dark and light grey boxes represent exons and introns, respectively. Introns have been scaled down as denoted for better presentation of the gene structures. Numbers at the scale bar indicate amino acids. (B) Schematic representation of the mouse SPIRE proteins. The *SPIRE1* gene encodes three different proteins, SPIRE1, SPIRE1-E9 and SPIRE1-E13. SPIRE1-E9 contains the alternatively spliced 14 amino acids spanning exon 9 located between the WH2 domains and the central MYO5 binding motif (GTBM). SPIRE1-E13 contains the alternatively spliced 58 amino acids spanning exon 13 located between the GTBM and the C-terminal membrane interaction module (SPIRE-box and FYVE-type zinc finger). The *SPIRE2* gene encodes for only one protein, SPIRE2. (C) RT-PCRs showing the presence of *SPIRE1* mRNA in general (total), *SPIRE1-E9* (E9) and *SPIRE1-E13* (E13) in mouse brain tissue (*left panel*). The *SPIRE2* mRNA was also present in brain (*middle panel*). A *SPIRE* mRNA containing both, exon 9 and exon 13 (*SPIRE1-E9E13*) was not detectable (*right panel*). For each RT-PCR cloned cDNA plasmids for each gene were used as positive controls and compared with the products from reactions using total RNA as template. Ethidium bromide stained agarose gels of the PCR products are shown. (D) Absolute quantification of mRNA copy numbers of *SPIRE1, SPIRE1-E9, SPIRE1-E13* and *SPIRE2* in mouse brain and small intestine as shown in a bar diagram. Each bar represents mean mRNA copy numbers of three independent experiments. *Error bars* represent SEM. (E) Myc-epitope (SPIRE1; *red*) and GFP (AcGFP1; SPIRE1-E9, SPIRE1-E13, SPIRE2; *green*) tagged SPIRE proteins are transiently co-expressed in HeLa cells. Deconvoluted images show the localization of SPIRE1, SPIRE1-E9 and SPIRE2 at vesicular membranes. SPIRE1-E13 is localized at mitochondria membranes as indicated by the colocalization with a mitochondrial stain (mitotracker orange; mitotracker; *red*). Colocalization is indicated by overlapping punctae (*yellow*). *Scale bar* represents 5 μm. At least 3 cells were recorded for each condition and the channel overlap of one representative cell is presented here. (F) The colocalization of tagged proteins and mitotracker, respectively, as described in (E) was quantified for the indicated conditions by determining its Pearson’s correlation coefficient (PCC) as shown in a bar diagram. Each bar represents the mean PCC value for at least 3 cells analyzed. *Error bars* represent SEM.

A distinct tissue expression of the *SPIRE1* and *SPIRE2* genes has been shown before (Schumacher, et al. 2004; Pleiser, et al. 2010; Pfender, et al. 2011). Besides oocytes which express high levels of both *SPIRE* genes (Pfender, et al. 2011), neuronal cells of the nervous system and epithelial cells of the digestive tract are major loci of *SPIRE1* and *SPIRE2* expression, respectively (Schumacher, et al. 2004; Pleiser, et al. 2010)(Fig. 6D). By employing absolute qPCR, with cloned cDNA vectors encoding the four mouse *SPIRE* isoforms as a standard, we quantified the absolute expression of each of the isoforms in brain and small intestine (Fig. 6D). The expression studies show that the amount of expression of *SPIRE1* and *SPIRE2* reverses between brain and the small intestine, with higher *SPIRE1* expression in brain and higher *SPIRE2* expression in the small intestine (Fig. 6D). In both tissues, the alternatively spliced isoforms *SPIRE1-E9* and *SPIRE1-E13* are only expressed in minor amounts never exceeding 9% for *SPIRE1-E9* and 5% for *SPIRE1-E13* of the total *SPIRE1* expression (Fig. 6D). In general, the absolute quantification showed that the mouse *SPIRE* genes are expressed at a very low level. By gene structure, tissue expression and subcellular localisation the single SPIRE protein as found in choanoflagellates has reached an increased complexity in mammals.

### Differential interaction of SPIRE1 and SPIRE2 with the RAB GTPases of the exocytic pathways

The complexity of cellular trafficking routes in regulating the communication of animal cells and tissues is most impressively shown by the segregation of the single RAB8 gene as found in the *M. brevicollis* genome into nineteen closely related human genes, including the RAB8, RAB27 and RAB3 branches (Klopper, et al. 2012; Rojas, et al. 2012; Varoqueaux and Fasshauer 2017). In order to understand whether the increased complexity of RAB8 related GTPases is also depicted in the complexity of mammalian SPIRE proteins, we have analysed the interaction of SPIRE1 and SPIRE2 with RAB8A, RAB27A and RAB3A (Fig. 7).

**Figure 7.**
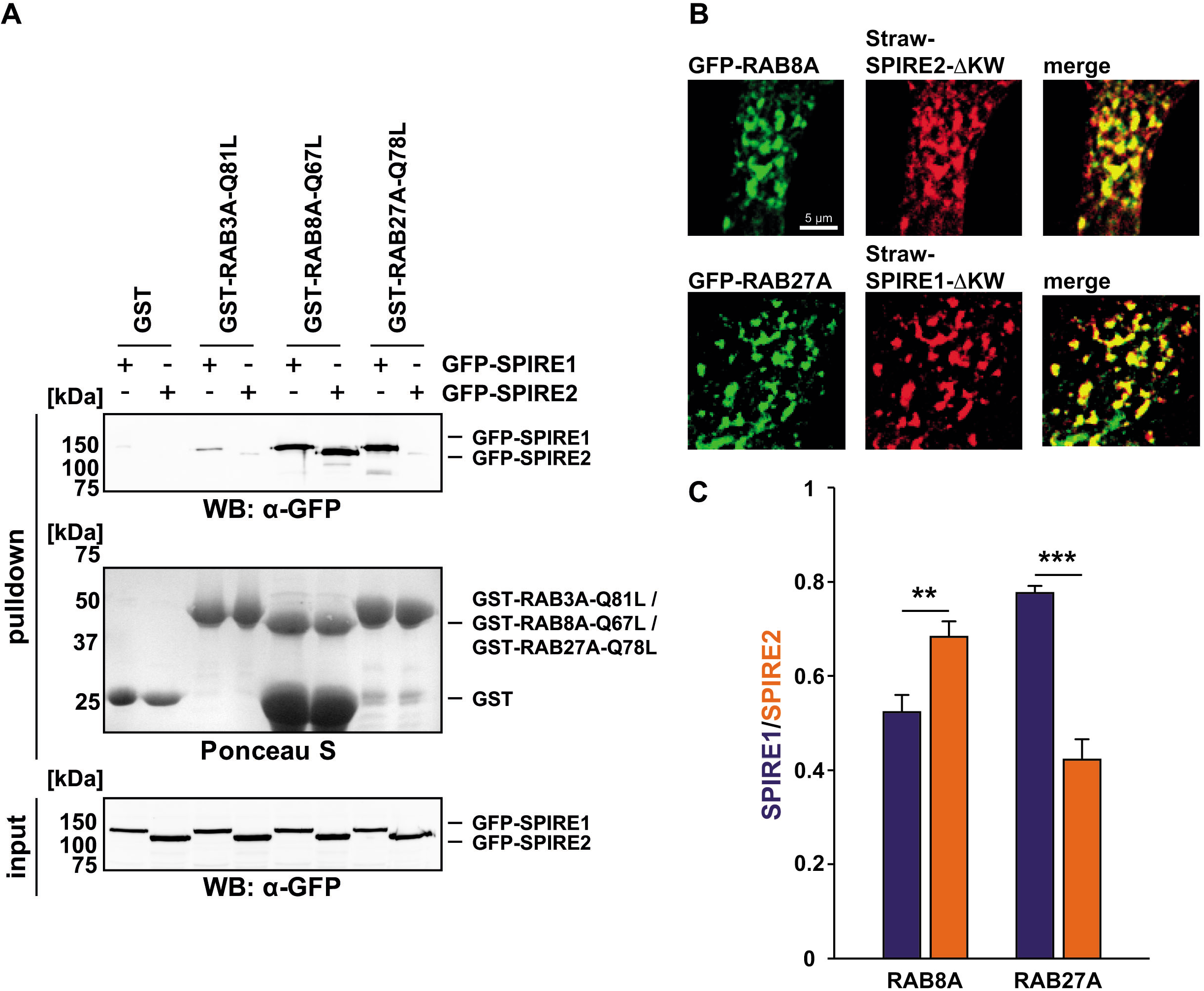
Vertebrate SPIRE1 and SPIRE2 proteins specifically interact with distinct RAB GTPases. (A) GST-pulldown assay with purified GTP-locked GST-RAB3A-Q81L, GST-RAB8A-Q67L and GST-RAB27A-Q78L and HEK293 cell lysates transiently over-expressing full-length AcGFP1-tagged Hs-SPIRE1/2 (GFP-SPIRE1 and GFP-SPIRE2). GST and GFP, respectively, were used as controls and Ponceau S staining shows equal amounts of GST-tagged proteins. N = 2 experimental repeats. (B) The localization of transiently co-expressed tagged RAB8A (eGFP; GFP-RAB8A; *green*) and SPIRE2-ΔKW (mStrawberry; Straw-SPIRE2-ΔKW; *red*) (*upper panel*) as well as of RAB27A (AcGFP1; GFP-RAB27A; *green*) and SPIRE1-ΔKW (mStrawberry; Straw-SPIRE1-ΔKW; *red*) (*lower panel*) was analyzed by fluorescence microscopy. Deconvoluted images indicate the localization of the proteins on vesicular structures and colocalization of RAB8A and SPIRE2 as well as of RAB27A and SPIRE1 (*merge*). *Scale bars* represent 5 μm. At least 5 cells were recorded for each condition and the cytoplasmic region of one representative cell is presented here. (C) The colocalization of tagged proteins as described in (A) was quantified for the indicated co-expressions of RAB8A and RAB27A with SPIRE1 and SPIRE2, respectively, by determining the Pearson’s correlation coefficient (PCC) as shown in a bar diagram. Each bar represents the mean PCC value for at least 5 cells analyzed. *Error bars* represent SEM. Statistical analysis was done using Student’s t-test to compare two co-expression conditions with a confidence interval of 95%. **p<0.01, ***p<0.001.

GTP-locked QL mutants of the three RAB GTPases were fused to GST and the corresponding proteins were purified from bacteria (GST-RAB3A-Q81L, GST-RAB8A-Q61L, GST-RAB27A-Q78L). GST-pulldown experiments employing GST-RAB proteins and AcGFP-tagged SPIRE1 and SPIRE2 transiently expressed in HEK293 cells revealed that both mammalian SPIRE proteins interact with comparable affinity with RAB8A (Fig. 7A). However, only the SPIRE1 protein strongly interacted with RAB27A (Fig. 7A). The interaction with RAB3A was found to be very weak as compared to the RAB8A and RAB27A interaction and might not be of physiological relevance (Fig. 7A). Colocalisation studies of transiently expressed mStrawberry-tagged SPIRE1 and SPIRE2 proteins with AcGFP-tagged RAB27A in HeLa cells revealed a much higher degree of colocalisation of RAB27A with SPIRE1 than SPIRE2, being consistent with the observed protein interaction study (Fig. 7B,C). Although we found a similar interaction of SPIRE1 and SPIRE2 with RAB8A-QL in GST-pulldown assays, colocalisation experiments in HeLa cells showed a preferential colocalisation of SPIRE2 with RAB8A as compared to SPIRE1 (Fig. 7C). The difference was, however, much less pronounced than in the case of RAB27A. In summary our expression and interaction studies show a specialisation of the vesicular SPIRE1 and SPIRE2 proteins. SPIRE2 is highly expressed in the digestive tract and has a preferential colocalisation with RAB8A. SPIRE1 is highly expressed in the brain and interacts and colocalizes preferentially with RAB27A. It is interesting to note that our preliminary data show that the mitochondrial *SPIRE1-E13* isoform is highest expressed in kidney, a tissue which requires extremely high amounts of energy to perform its blood filtration function.

## Discussion

Vesicle transport processes in animal cells can be described by the highways and local roads model, which addresses a cooperative transport along microtubule and actin filament tracks (Hammer and Sellers 2011; Hume and Seabra 2011). In plant cells actin and microtubule transport are separated and in fungi actin cables are the major transport routes (Bretscher 2003; Geitmann and Nebenfuhr 2015)(Fig. 8). We here show that in Holozoa, in the single cell ancestor of the animal kingdom, a new transport mechanism emerged, which is molecularly presented by the FMN/SPIRE/MYO5 complex and provides an additional degree of freedom in cargo transport by organising local actin/myosin networks directly at vesicle membranes (Fig. 8A, B). Based on the function of the FMN/SPIRE/MYO5 cooperation in vertebrate oocytes and melanocytes, we propose a model in which these local actin networks might in general bridge the gap between microtubule tracks, detached vesicles and the cellular actin cytoskeleton (Fig. 8A, B). The enhanced performance in vesicle motility by SPIRE/FMN generated actin networks may contribute to the complexity of cellular communications in animals. Mouse genetics showed that this specifically holds true for neuronal communication in the brain, by showing a function of SPIRE1 and FMN2 in fear learning (Peleg, et al. 2010; Pleiser, et al. 2014; Agis-Balboa, et al. 2017). In analogy, humans with a homozygous FMN2 mutation suffer from intellectual disability (Law, et al. 2014).

**Figure 8.**
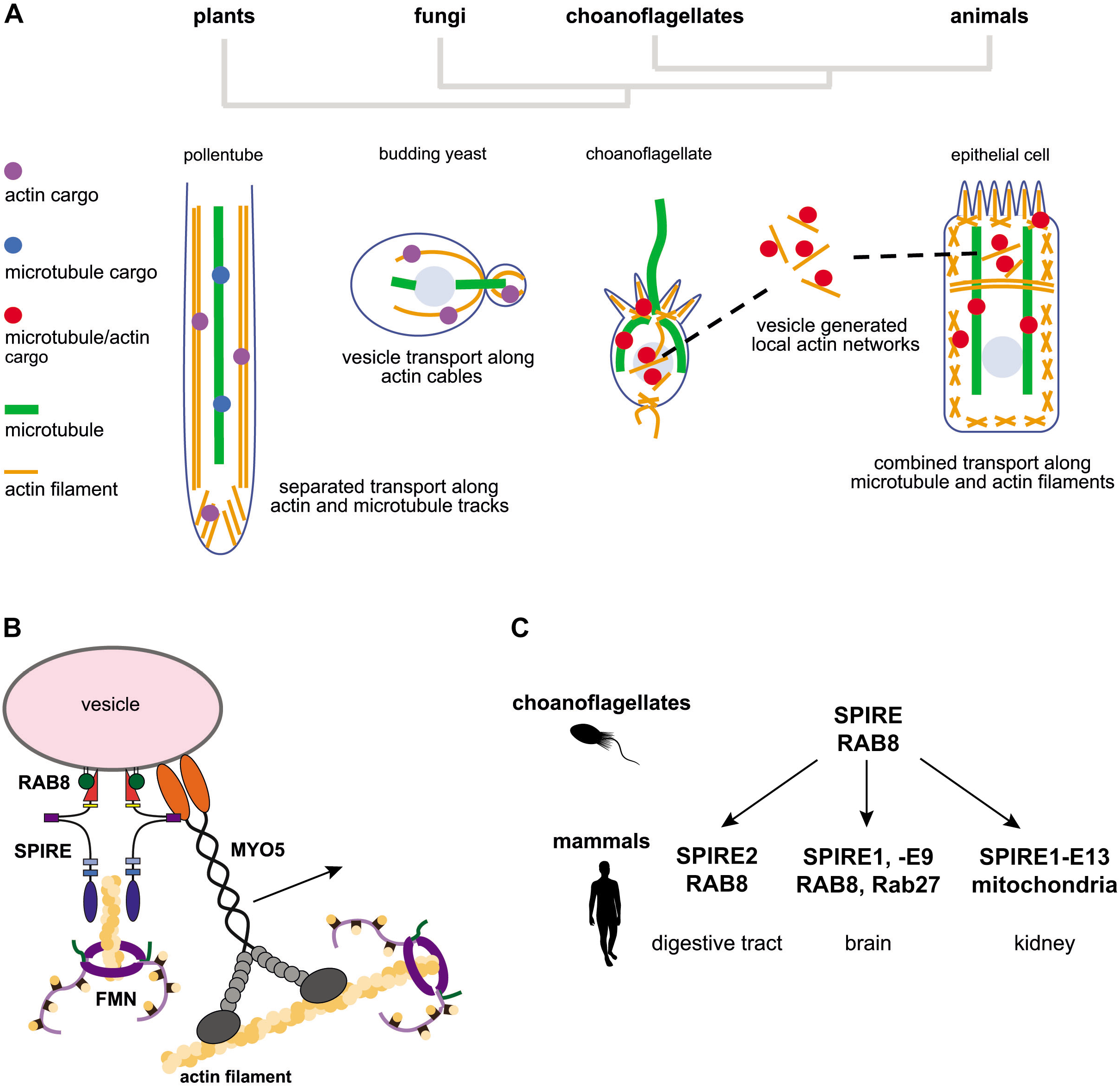
The evolution of local actin/myosin networks at vesicle membranes. (A) Schematic presentations of vesicle transport along actin and microtubule tracks in plants, yeast, choanoflagellates and mammalian cells. (B) Model of the choanoflagellate RAB8/SPIRE/MYO5/FMN protein complex at vesicle membranes. (C) The mammalian genomes encode two *SPIRE* genes, which separated SPIRE protein function starting from its original SPIRE/RAB8 complex in terms of tissue expression, RAB GTPase binding and mitochondria localisation.

Our genome analysis and interaction data show that the FMN/SPIRE/MYO5 complex first comes into existence in holozoan sister taxa to metazoans, which represent the closest single cell relatives of animals. The cytoskeleton of choanoflagellates consists of cytoplasmic microtubule tracks just like vertebrate cells and also cortical actin structures, such as the actin-filled villi of the collar and filopodia extending from the basal pole of the cell (Hoffmeyer and Burkhardt 2016; Brunet and King 2017; Booth, et al. 2018). Interestingly there are also actin filaments originating in the cell body, which coalesce at the base of the collar (Booth, et al. 2018). These actin filaments may be related to actin cables of plant and yeast cells, which function as vesicle and organelle transport tracks. The transport mechanisms in Choanoflagellida and Ichthyosporea still remain to be analysed. The structural array of the choanoflagellate cytoskeleton, however, very much resembles that of epithelial cells and may therefore also have combined actin/microtubule transport processes.

We have identified the choanoflagellate SPIRE and the mammalian SPIRE1 and SPIRE2 proteins as binding partners of the RAB8 GTPase (Fig. 3, Fig. 7). In mammalian intestinal cells transport regulated by RAB8 and the actin motor protein MYO5B functions in apical protein localisation (Sato, et al. 2007; Vogel, et al. 2015). In analogy to the RAB8/MYO5B function in the mammalian intestine, choanoflagellate RAB8 and MYO5 may be involved in transport processes towards the choanoflagellate collar. It seems reasonable to assume that similar interactions will be found in ichthyosporean species. The preferential interaction of mammalian SPIRE2 with RAB8A in combination with the high expression of SPIRE2 in epithelial cells of the digestive tract, suggests a function of SPIRE2 in apical transport processes of epithelial cells.

The genome duplications at the origin of the vertebrates increased the complexity of SPIRE function. An alternatively spliced exon targets the vertebrate SPIRE1-E13 towards mitochondria (Manor, et al. 2015)(Fig. 6) and the vesicular SPIRE1 protein gained specificity for the interaction with the RAB27 GTPase (Fig. 7, Fig. 8). RAB27 regulates secretory processes and cellular communication (Fukuda 2013). By sequentially recruiting effector proteins to secretory vesicles, RAB27 escorts the secretion pathway from the microtubule tracks to the actin cytoskeleton and finally to membrane fusion. Two distinct RAB27 effectors along this pathway directly interact with the actin cytoskeleton, the SPIRE proteins and the melanophilin (MLPH)/MYRIP proteins (Alzahofi et al., under revision)(Fukuda and Kuroda 2002; Quinlan, et al. 2005). We have shown before that SPIRE and MLPH share considerable similarity in both their RAB27 interaction domain and their MYO5 interaction motif (Alzahofi et al., under revision)(Pylypenko, et al. 2016). The proteins, however, differ in the interaction with the actin cytoskeleton. Whereas SPIRE proteins catalyse actin filament generation (Quinlan, et al. 2005; Dahlgaard, et al. 2007; Quinlan, et al. 2007; Pfender, et al. 2011), MLPH and MYRIP are actin filament binding proteins (Fukuda and Kuroda 2002; Ramalho, et al. 2009). *In vitro* assays show that MLPH increases the number of processively moving MYO5A motors and nearly doubles the run length of the motors (Sckolnick, et al. 2013). It has been suggested that MLPH acts as a tether that links the MYO5A motor to the actin track. In melanocytes RAB27A, SPIRE1, FMN1, MYO5A and MLPH cooperate in cytoplasmic melanosome dispersion (Alzahofi et al., under revision) and it would be interesting to know if cooperation with MLPH or MYRIP is a general feature of SPIRE proteins throughout evolution. The very low conservation of MYRIP sequences outside the vertebrate family makes it very difficult to resolve the evolution of MYRIP/MLPH function. By now we were not able to identify a MYRIP/MLPH homolog in choanoflagellates. Further genome studies will be required to address a possible coevolution of SPIRE and MYRIP/MLPH function.

The combined genomic/molecular cell biology study performed here has provided very valuable information on the evolution of actin/myosin networks at vesicle membranes in the animal kingdom. The characterisation of SPIRE proteins as RAB8/RAB27 effectors indicates that the new transport function specifically evolved to facilitate exocytic transport, which is the basis of cellular communication. Our data are important to further explore the yet poorly understood functions of SPIRE proteins in mammalian tissues.

## Materials and methods

### Identification and annotation of SPIRE genes

*SPIRE* genes have been identified and annotated using the same approach, with which thousands of tubulins, myosins and dynein heavy chains were annotated and which has been described in exhaustive detail there (Findeisen, et al. 2014; Kollmar 2016; Kollmar and Muhlhausen 2017). Shortly, starting with the protein sequences of human and *Drosophila melanogaster* SPIRE, further homologs were identified in TBLASTN searches of available genome and transcriptome assemblies. Hits in transcriptome assemblies were translated in the respective reading frame. Hits in genome assemblies served to identify gene regions, and the respective genomic regions were submitted to AUGUSTUS (Stanke, et al. 2004) to obtain gene predictions. These gene predictions were subsequently manually corrected and refined. In total, 321 SPIRE genes were assembled. All sequence related data (domain predictions, gene structures, sequences) and references to genome sequencing centres are available at CyMoBase (www.cymobase.org) (Odronitz and Kollmar 2006).

### Generating the multiple sequence alignment

SPIRE proteins contain multiple short sequence motifs such as WH2, GTBM and SB motifs (Welz and Kerkhoff 2017), which are connected by highly divergent linker sequences. Therefore, generating multiple sequence alignments with software such as Muscle, MAFFT or ClustalW results in highly fragmented alignments and the four WH2 domains are often not aligned in correct order and GTBM motifs not recognized at all. Thus, we generated a preliminary alignment using ClustalW (Chenna, et al. 2003), which we manually refined extensively. Based on this alignment, we further corrected gene predictions by removing wrongly predicted sequence regions and by filling gaps. In case SPIRE sequence gaps remained because of genome assembly gaps, we maintained the integrity of exons 5’ and 3’ of gaps. The SPIRE sequence alignment is available from CyMoBase (www.cymobase.org) (Odronitz and Kollmar 2006) and as Supplementary Data S1.

### Computing and visualising phylogenetic trees

The alignment was treated with CD-Hit v.4.5.4 (Li and Godzik 2006) applying a similarity threshold of 90% to generate a dataset with less redundancy. The resulting dataset contained 217 sequences and 2,447 alignment positions. IQ-TREE v.1.6.beta3 ModelFinder (Nguyen, et al. 2015) was used to determine the most appropriate amino acid substitution model. ModelFinder identified the JTT+F+R8 (Jones, et al. 1992) to be the best model under the Bayesian Information Criterion (BIC). Phylogenetic trees were generated using the Maximum likelihood method. Maximum likelihood (ML) analyses with estimated proportion of invariable sites and bootstrapping (1,000 replicates) were performed with FastTree v.2.1.10 SSE3 (Price, et al. 2010) using the settings for increased accuracy (`–spr 4 –mlacc 2 –slownni`) as described in the documentation. IQ-TREE-generated trees showed identical topology of almost all branches and slightly varying bootstrap values (1,000 replicates) at some of the less supported nodes. Phylogenetic trees were visualized with FigTree (http://tree.bio.ed.ac.uk/software/figtree/).

### WebLogo sequence alignment

WebLogos (Schneider and Stephens 1990; Crooks, et al. 2004) were generated online using the WebLogo server (weblogo.berkeley.edu) from the respective parts of chosen vertebrate FMN1/2 sequences to depict conserved amino acids within the FMN-FSI motif and of chosen vertebrate SPIRE1/2 sequences to depict conserved amino acids within the SPIRE-GTBM.

### Gene synthesis

A full-length coding region of the *Mb-SPIRE* cDNA, a coding region of the *Mb-MYO5* globular tail domain cDNA, a coding region of the *Mb-FMN* C-terminal cDNA sequences (FH2-FSI) and a full-length coding region of the *Mb-RAB8* cDNA were generated by gene synthesis (Eurofins Genomics Germany, Ebersberg, Germany) and were codon-optimized for expression in *E.coli* bacterial cells. Synthesized cDNA sequences were subsequently used as templates for further subcloning of respective cDNA fragments to allow for bacterial and eukaryotic protein expression.

### Expression vector cloning

Expression vectors were generated by standard cloning techniques using AccuPrime Pfx DNA polymerase (Thermo Fisher, Waltham, MA, USA), restriction endonucleases and T4 DNA ligase (both New England Biolabs, Frankfurt am Main, Germany). Point mutants were generated using the In-Fusion HD cloning kit (TakaraBio/Clontech, Saint-Germain-en-Laye, France). DNA sequencing was carried out by LGC Genomics (Berlin, Germany). Table 1 shows details of vectors used in this study.

**Table 1.**
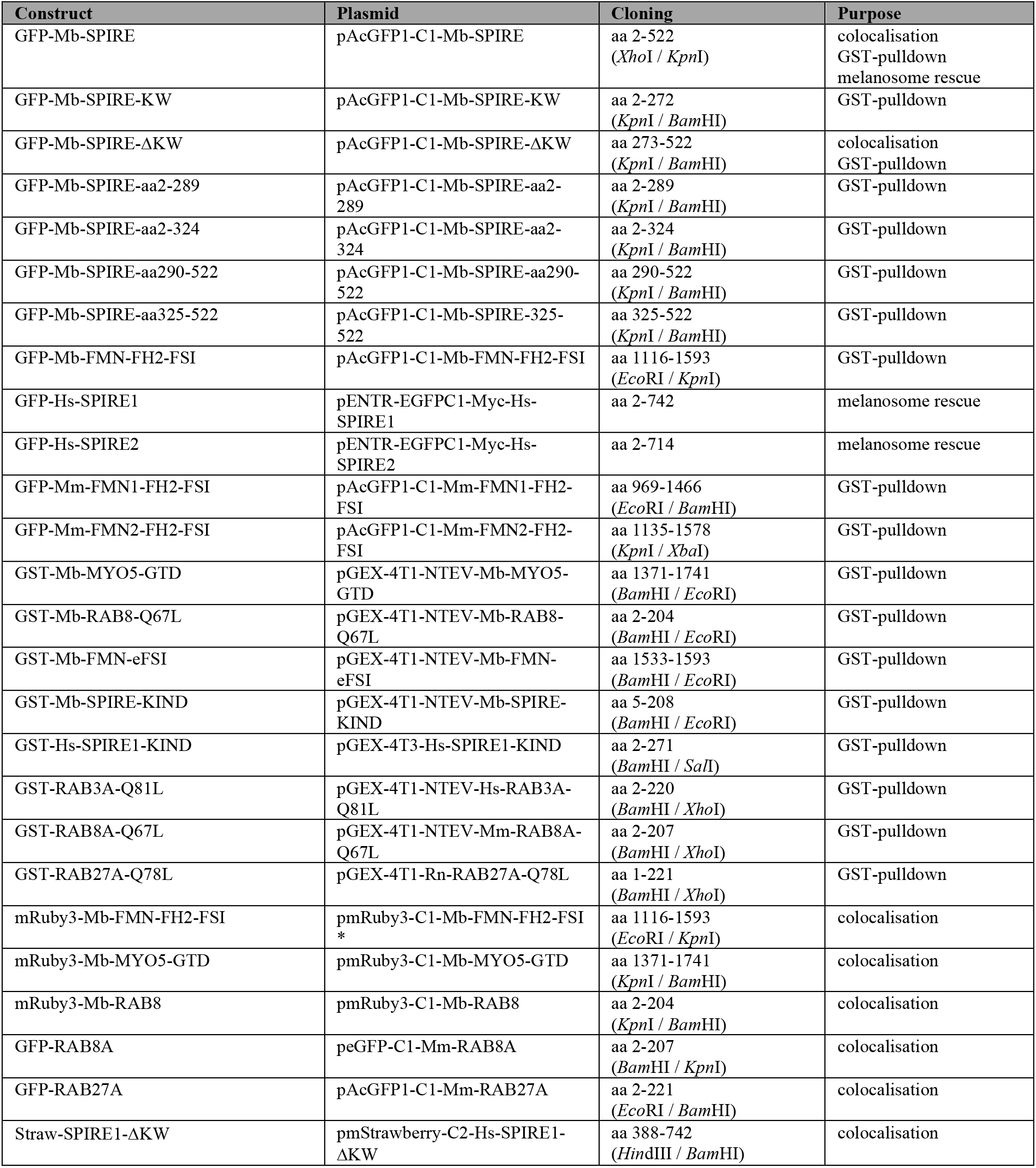
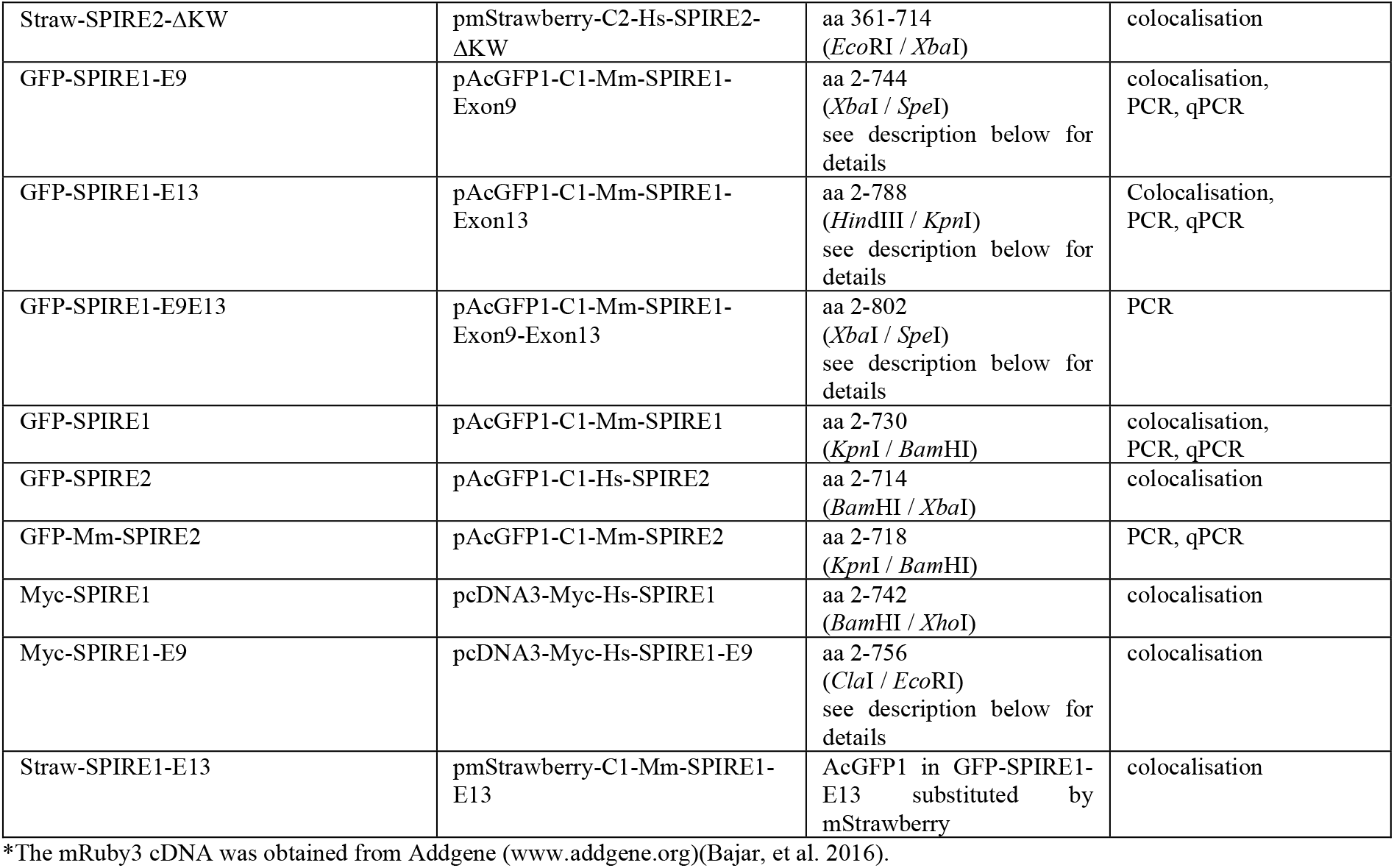
Prokaryotic and eukaryotic protein expression vectors used in this study.

### Cloning of SPIRE1-E9, SPIRE1-E13 and SPIRE1-E9E13

Expression vectors encoding human and mouse SPIRE1-E9 proteins (pcDNA3-Myc-Hs-SPIRE1-E9 and pAcGFP1-C1-Mm-SPIRE1-E9), respectively, were generated using PCR amplification of two DNA fragments and subsequent assembly PCR (using Pfu DNA polymerase from Promega, Mannheim, Germany) with the artificially inserted exon 9 sequence as region for annealing. The obtained assembly PCR SPIRE1 fragments including exon 9 were inserted into existing SPIRE1 expression vectors by using specific restriction endonuclease sites (see also Tab. 1). An expression vector encoding the mouse SPIRE1-E13 protein (pAcGFP1-C1-Mm-SPIRE1-E13) was generated using PCR amplification of two DNA fragments from mouse cerebellum cDNA template. 3’ primer of PCR1 and 5’ primer of PCR2 were chosen to anneal specifically to exon 13 encoded cDNA sequences. Both PCR fragments were subsequently used for assembly PCR with exon 13 encoded sequences as region for annealing. The obtained assembly PCR SPIRE1 fragment including exon 13 was inserted into an existing SPIRE1 expression vector by using specific restriction endonuclease sites (see also Tab. 1). An expression vector encoding the physiologically not expressed mouse SPIRE1-E9E13 protein carrying both alternatively spliced exon 9 and 13 (pAcGFP1-C1-Mm-SPIRE1-E9E13) was generated using PCR amplification of two DNA fragments from pAcGFP1-C1-Mm-SPIRE1-E13 plasmid DNA template (see above) and subsequent assembly PCR with the artificially inserted exon 9 sequence as region for annealing. The obtained assembly PCR SPIRE1-E9E13 fragment including exon 9 (from primer) and exon 13 (from template) was inserted into an existing SPIRE1 expression vector by using specific restriction endonuclease sites (see also Tab. 1).

### Recombinant protein expression and purification

Recombinant GST-Mb-MYO5-GTD, GST-Mb-FMN-eFSI, GST-Mb-RAB8-Q67L, GST-Mb-SPIRE-KIND, GST-Hs-SPIRE1-KIND, GST-Hs-RAB3A-Q81L, GST-Mm-RAB8A-Q67L and GST-Rn-RAB27A-Q78L proteins were expressed in *Escherichia coli* Rosetta bacterial cells (Merck Millipore, Novagen, Darmstadt, Germany). Bacteria were cultured in LB medium (100 mg/l ampicillin, 34 mg/l chloramphenicol) at 37°C until an A600 nm of OD 0.6 - 0.8. Protein expression was induced by 0.2 mM Isopropyl-²-D-thiogalactopyranoside (IPTG; Sigma-Aldrich, Taufkirchen, Germany) and continued at 16 - 20°C for 18 - 20 h. Bacteria were harvested and lysed by ultra-sonication. Soluble proteins were purified using GSH-Sepharose 4B (GE Healthcare Life Sciences) and size exclusion chromatography (High Load 16/60 Superdex 200; GE Healthcare Life Sciences). Proteins were concentrated by ultrafiltration using Amicon Ultra centrifugal filters (Merck Millipore) with respective molecular weight cut offs. The final protein purity was estimated by SDS-PAGE and Coomassie staining. Purification of GST-Mm-RAB8A-Q67L was challenging because of low protein solubility and aggregation into inclusion bodies. Therefore, volume of bacterial cultures was increased to 5 l to get a reasonable amount of purified protein.

### Cell culture

HEK293 and HeLa cells (both from ATCC, Manassas, Virginia, USA) were cultured in Dulbecco’s Modified Eagle’s Medium (DMEM; Thermo Fisher) supplemented with 10% (v/v) fetal calf serum (FCSIII; GE Healthcare Life Sciences, HyClone), 2 mM L-glutamine (Thermo Fisher), penicillin (100 units/ml; Thermo Fisher) and streptomycin (100 μg/ml; Thermo Fisher) at 37°C, 5% CO_2_, 95% humidity and were passaged regularly at 80% confluency. Transfections with plasmid DNA were performed using Lipofectamine2000 reagent (Thermo Fisher) according to manufacturer’s recommendation. Cultures of immortal melan-a melanocytes (available from the Wellcome Trust Functional Genomics Cell Bank at St George’s University of London UK SW17 0RE) were maintained as described previously (Hume, et al. 2001).

### siRNA transfection of melanocytes

siRNA transfection of melanocytes using oligofectamine was performed as previously described (Hume, et al. 2007). NT, RAB27A specific siRNA were as previously described (Robinson, et al. 2017). The sequence of siRNA oligonucleotides was GGACGACAUUCGGUGCAAA for mouse SPIRE1 and CAAAGAACACUGCACGAGA for mouse SPIRE2. All siRNA oligonucleotides were from Sigma Genosys UK.

### Generation of adenovirus vectors for protein expression in mammalian cells

Adenovirus expression vectors were generated as previously described (Hume, et al. 2006).

### GST-pulldown protein interaction assays

For GST-pulldowns HEK293 cells were transfected with expression vectors encoding GFP-tagged Mb-SPIRE full-length proteins and respective deletion mutants, Hs-SPIRE1, Hs-SPIRE2 and C-terminal fragments of Mb-FMN, Mm-FMN1 and Mm-FMN2. GFP alone was used as a control. 48 h post transfection, cells were lysed in lysis buffer (25 mM Tris-HCl pH 7.4, 150 mM NaCl, 5 mM MgCl_2_, 10% (v/v) glycerol, 0.1% (v/v) Nonidet P-40, 1 mM PMSF, protease inhibitor cocktail) and centrifuged at 20,000 x g, 4°C, 20 min to remove insoluble debris. For GST-pulldown assays 65 μg GST-Mb-MYO5-GTD, 32 μg GST-Mb-FMN-eFSI, 47 μg GST-Mb-RAB8-Q67L, 50 μg GST-Hs-RAB3A-Q81L, 50 μg GST-Mm-RAB8A-Q67L, 50 μg GST-Rn-RAB27A-Q78L, 50 μg GST-Mb-SPIRE-KIND and 50 μg GST-Hs-SPIRE1-KIND protein, respectively, and 25 μg GST protein as control was coupled to GSH-Sepharose 4B beads (1:1 suspension) for 1 h, 4°C on a rotating wheel. Beads were washed twice with pulldown buffer (25 mM Tris-HCl pH 7.4, 150 mM NaCl, 5 mM MgCl_2_, 10% (v/v) glycerol, 0.1% (v/v) Nonidet P-40) and subsequently incubated with the cell lysates for 2 h at 4°C on a rotating wheel.

Beads were washed four times with pulldown buffer and bound proteins were eluted with 1x Laemmli buffer, denatured at 95°C for 10 min and then analyzed by immunoblotting.

### Immunoblotting

Immunoblotting was performed as described previously (Pylypenko, et al. 2016) using anti-GFP (Living Colors A.v. peptide antibody, rabbit polyclonal, 1 mg/ml; TakaraBio/Clontech) primary antibody and horseradish peroxidase linked anti-rabbit IgG (from donkey) secondary antibody (1:5000, GE Healthcare Life Sciences). Signal was detected by chemiluminescence (Luminata Forte Western HRP substrate; Merck Millipore) and recorded with an Image Quant LAS4000 system (GE Healthcare Life Sciences).

### Immunostaining

HeLa cells were seeded on microscope cover glasses and transfected to transiently express fluorescently-tagged Mb-SPIRE, Mb-FMN, Mb-RAB8, Mb-MYO5, Mm-SPIRE1-E9, Mm-SPIRE1-E13, Hs-SPIRE2, Mm-RAB8A, Rn-RAB27A, Hs-SPIRE1-ΔKW and Hs-SPIRE2-ΔKW proteins and Myc-epitope tagged Hs-SPIRE1 protein, respectively. Cells were fixed with paraformaldehyde (3.7% in 1x PBS) for 20 min at 4°C and subsequently permeabilized using 0.2% Triton X-100 (in 1x PBS) for 3.5 min, room temperature. Respective cells were incubated with anti-c-Myc antibody (9E10, mouse monoclonal, 2 mg/ml; Santa Cruz Biotechnology, Dallas, TX, USA) for 1 h at room temperature, and conjugated anti-mouse secondary antibodies (Cy5; from donkey, 3.25 mg/ ml; Dianova, Hamburg, Germany) for 1 h at room temperature avoiding exposure to light. Finally, cells were mounted on microscope slides with Mowiol, dried at room temperature in the dark and stored at 4°C.

### Fluorescence microscopy

Fixed cells were analyzed with a Leica AF6000LX fluorescence microscope, equipped with a Leica HCX PL APO 63x/1.3 GLYC objective and a Leica DFC7000GT CCD camera (16 bit, pixel size: 4.54 × 4.54 μm, 1920 × 1440 pixels, 2 × 2 binning mode). 3D stacks were recorded and processed with the Leica deconvolution software module. Images were recorded using the Leica LASX software and further processed with Adobe Photoshop and subsequently assembled with Adobe Illustrator.

### Mitochondrial staining for fluorescence live cell imaging

In order to fluorescently label mitochondrial membranes in living HeLa cells the cell permeable compound MitoTracker Orange (Thermo Fisher) was used. HeLa cells were seeded in glass-bottom tissue culture dishes (WillCo-Dish, 40 mm; WillCo Wells B.V., Amsterdam, The Netherlands) 20 h prior to the experiment. On the next day cells were washed in Opti-MEM (Thermo Fisher) and incubated with 2 ml of 5 nM MitoTracker Orange diluted in Opti-MEM for 20 min at 37°C, 5% CO_2_. Cells were washed again with Opti-MEM to get rid of excess staining solution and covered with Opti-MEM for subsequent fluorescence live cell imaging. Here the Leica AF6000LX fluorescence imaging system was used with the setup as described above. Single images were recorded using the Leica LASX software and further processed with Adobe Photoshop and subsequently assembled with Adobe Illustrator.

### Colocalization analysis

The extent of colocalization of SPIRE1 with SPIRE1-E9, SPIRE1-E13 and SPIRE2, respectively, at vesicle surfaces and of SPIRE1-E13 with mitochondrial membranes as well as of SPIRE1 and SPIRE2 with RAB8A and RAB27A at vesicular membranes was analyzed using the ImageJ (V2.0.0) plug-in Coloc2. Here, the colocalization rate is indicated by the Pearson’s Correlation Coefficient (PCC) as a statistical measure to unravel a linear correlation between the intensity of different fluorescent signals. A PCC value of 1 indicates a perfect colocalization, 0 indicates a random colocalization and a PCC value of −1 indicates a mutually exclusive localization of the analyzed signals. To take the noise of each image into account and to gain an objective evaluation of PCC significance, a Costes significance test was performed. To do so, the pixels in one image were scrambled randomly and the correlation with the other (unscrambled) image was measured. Significance regarding correlation was observed when at least 95% of randomized images show a PCC less than that of the original image, meaning that the probability for the measured correlation of two colors is significantly greater than the correlation of random overlap (Costes, et al. 2004; Pompey, et al. 2013).

### Qualitative PCR and quantitative real-time PCR

Male C57BL/6N mice were ordered from Charles River Laboratories (Sulzfeld, Germany) at an age of 6 weeks. Immediately after arrival mice were killed by cervical dislocation and organs of interest were isolated. Organs were weighted and stored in an adequate amount of *RNA later* (Qiagen, Hilden, Germany) at 4°C, which inhibits RNA degradation. For total RNA preparations of organs, the RNeasy Midi Kit (Qiagen) was used according to manufacturer’s recommendation. Disruption and homogenization of organs was done with the TissueRuptor (Qiagen). In order to determine amount and purity of isolated total RNA, a spectrophotometric analysis was done. Sample integrity of isolated RNA was measured with the QIAxcel (Qiagen) using the QIAxcel RNA QC Kit v2.0 (Qiagen) according to protocol. Subsequently cDNA was generated employing the QuantiNova Reverse Transcription Kit (Qiagen) according to recommendations. The reverse transcription kit includes a genomic DNA removal step. An internal control RNA was employed to verify successful reverse transcription. Standard qualitative PCRs to check for the general abundance of distinct *SPIRE1* splice variants were performed employing cDNAs from mouse brain tissue and Q5 High-Fidelity DNA polymerase (New England Biolabs). Expression vectors encoding mouse *SPIRE1, SPIRE1-E9, SPIRE1-E13, SPIRE1-E9E13* and *SPIRE2* cDNAs were used as positive controls.

The absolute quantification by quantitative real-time PCR (qPCR) was performed with the Rotor-Gene Q thermocycler (Qiagen) and the QuantiNova SYBR Green PCR Kit. Each reaction was set up in triplicates and C_T_ values were calculated by the Qiagen Rotor-Gene Q Series Software. For absolute quantification of sample mRNA copy numbers expression vectors encoding mouse *SPIRE1, SPIRE1-E9, SPIRE1-E13* and *SPIRE2* cDNAs were used to create a standard curve with known copy number concentrations. Therefore, plasmid DNA was linearized by restriction digest and then serially diluted in water. The copy number of standard DNA molecules was then determined by the following formula (from the script “Critical Factors for Successful Real Time PCR”, Qiagen):

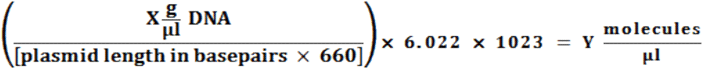

CT values of the samples were compared to those of the standard curve with known concentrations to calculate the sample mRNA copy numbers. The primer pairs used for both qualitative and quantitative PCRs are shown in table 2. The primer pair for “SPIRE1” amplifies all three SPIRE1 splice variants whereas the primer pairs for “SPIRE1-E9” and “SPIRE1-E13” are specific for the distinct splice variant. To determine the exact copy number of SPIRE1 mRNA, the calculated copy numbers of SPIRE1-E9 and SPIRE1-E13 were subtracted from the calculated total SPIRE1 copy number.

**Table 2.**
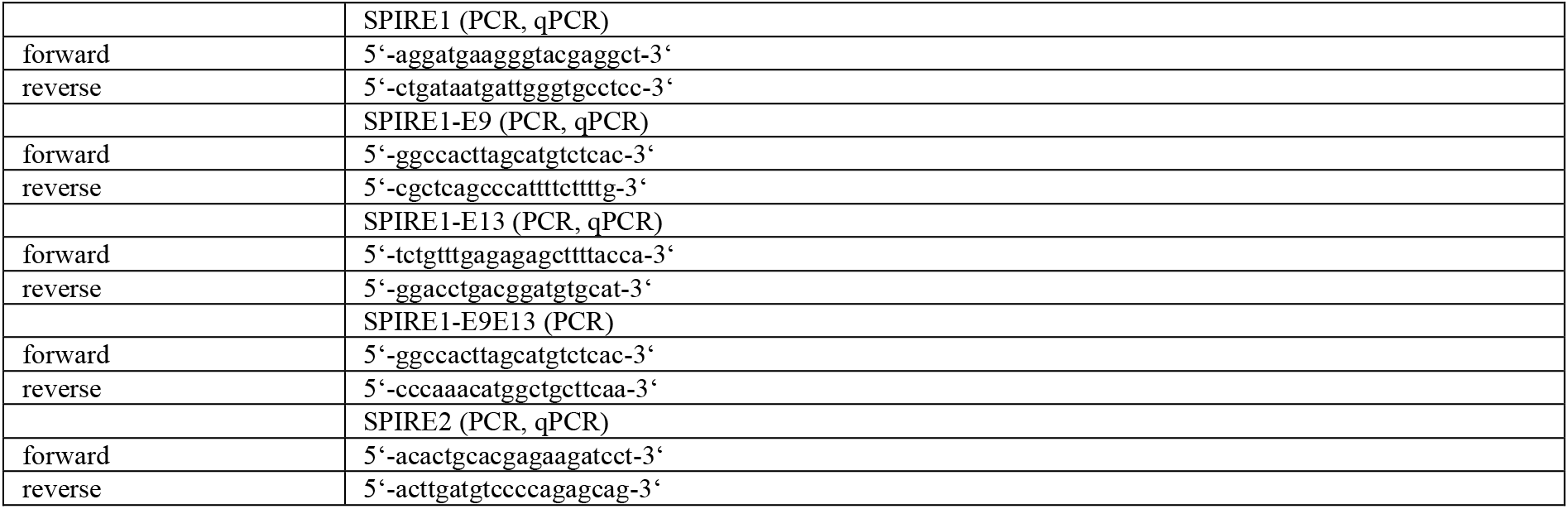
Primer pairs used for qualitative and quantitative PCRs of *SPIRE* variants.

### PCR primer design

Primer pairs for qPCR were designed to amplify exon specific sequences assisted the web-based tool Primer3 (http://primer3.ut.ee). Specificity of primer pairs was checked by performing a Primer-BLAST search (www.ncbi.nlm.nih.gov/blast). Primer oligonucleotides consist of 18 to 30 nucleotides and have a GC content of 40-60%. qPCR primer pairs were chosen to amplify less than 150 base pairs and flank a region that contains at least one intron. All primers were synthesized by Sigma-Aldrich and purified by HPLC. Lyophilized primers were resuspended in DEPC-H2O and stored at −20°C. Prior to use in qPCR all primers were tested to amplify a specific band in PCR and agarose gel electrophoresis.

## Acknowledgements

MK would like to thank Prof. Christian Griesinger for his continuous generous support. This project has been funded by grants SPP 1464: KO 2251/13-1 (to MK) and SPP 1464: KE 447/10-1, −2 and KE 447/18-1 (to EK) of the Deutsche Forschungsgemeinschaft (DFG). Felix Straub is a member of the DFG graduate school GRK 2174 (neurobiology of emotion dysfunctions).

